# A GPVI-platelet-neutrophil-NET axis drives systemic sclerosis

**DOI:** 10.1101/2025.01.21.634123

**Authors:** Roxane Darbousset, Gonzalo Villanueva-Martin, Julia Kuehn, Stefano Navarro, Spoorthi Balu, Dhruv Miglani, Eilish Dillon, Caroline Fromson, Leetah Senkpeil, Mehreen Elahee, Pierre-Andre Jarrot, Sydney B. Montesi, Deepak A. Rao, Jerry Ware, Denisa D. Wagner, Andreea M. Bujor, Bernhard Nieswandt, Maria Gutierrez-Arcelus, Peter A. Nigrovic

## Abstract

Systemic sclerosis (SSc) is immune-mediate inflammatory disease characterized by progressive tissue fibrosis. We observed that circulating neutrophils from patients with diffuse SSc exhibit an activated phenotype, a finding echoed in blood and skin transcriptomes. Neutrophil depletion abrogated experimental SSc induced by cutaneous injection of hypochlorous acid (HOCl) or bleomycin (BLM), and adoptive transfer of HOCl and BLM neutrophils induced skin and lung fibrosis in healthy mice, establishing neutrophils as necessary and sufficient for fibrosis. We noted that SSc patients exhibited platelet activation, a phenotype that preceded neutrophil activation in mice, suggesting an upstream role. Indeed, platelet depletion abrogated neutrophil activation and tissue fibrosis, and exposure to HOCl or BLM platelets conferred upon wild-type neutrophils the capacity to induce skin and lung fibrosis via neutrophil extracellular traps (NETs). Genetic and therapeutic blockade of the platelet collagen receptor GPVI attenuated platelet and neutrophil activation, reduced circulating NETs, and protected animals from skin and lung fibrosis. These findings identify the GPVI-platelet-neutrophil-NET as a new source of therapeutic targets in SSc.

## INTRODUCTION

Systemic sclerosis (SSc), also termed scleroderma, is a chronic immune-mediated inflammatory disease characterized by vascular abnormalities, immune system dysregulation, and widespread deposition of excess extracellular matrix (*1–4*). Fibrosis affects the skin but can also involve the lungs, gastrointestinal tract, and heart, especially in diffuse SSc (dSSc), the most aggressive form of the disease, leading to high morbidity and mortality (*3, 5, 6*). Therapeutic options for SSc remain limited.

Inflammation can precede and accompany fibrosis in SSc (*7–10*). Neutrophils, the cellular hallmark of acute inflammation, are associated with poor prognosis in SSc when abundant in blood or bronchoalveolar (BAL) fluid (*11–13*). Yet evidence for neutrophil activation in SSc has varied between studies, and the role of these cells as pathogenic effectors is unresolved (*14–18*).

Endothelial injury occurs early in SSc and is generally regarded as the primary tissue lesion, as suggested by clinical and histological studies and by the observation that environmental agents linked to SSc-like syndromes are commonly endothelial toxins (*19, 20*). Multiple mechanisms connect vascular injury to tissue fibrosis, including immune cell activation, hypoxia, thrombosis, and endothelial-to-mesenchymal transition (*21, 22*). In this context, platelets are of particular interest. Platelets from SSc patients exhibit evidence of activation, with enhanced expression of surface markers such as P-selectin (*23–25*). Some authors have identified unusual susceptibility of SSc platelets to activation by collagen, a protein exposed when the endothelial lining is disrupted (*19, 26–29*). Patients with SSc exhibit an abundance of circulating platelet-derived microparticles capable of activating neutrophils and potentially other cells, and adoptive transfer of these microparticles mediates endothelial injury in lung (*17, 30*). Nevertheless, how platelets become activated and whether platelets and neutrophils play key roles in SSc *in vivo* remains to be established.

Here we studied circulating neutrophils and platelets in patients with dSSc and tested the contribution of neutrophils, neutrophil extracellular traps (NETs), and platelets to fibrosis in murine models commonly employed to model SSc. We establish that both neutrophils and platelets play essential roles in the pathogenesis of experimental SSc, in a cascade initiated through the platelet collagen receptor glycoprotein VI (GPVI) and ending with platelet-induced NET formation. Murine disease can be attenuated by intervention at any point in this cascade, defining potential new therapeutic targets in this challenging disease.

## RESULTS

### Neutrophils are activated in diffuse systemic sclerosis

We began by characterizing neutrophils from the peripheral blood of patients with dSSc compared with healthy controls (Supplemental Table 1 and 2 for demographic and phenotypic information). In dSSc patients, the proportion of neutrophils among all leukocytes and the abundance of these cells in circulation were both increased, as was the neutrophil-to-lymphocyte ratio (Figure 1 A-C). Circulating neutrophils expressed more total and active-conformation Mac-1 integrin and trended lower in L-selectin, all features of activation (Supplementary Figure 1 A for gating strategy; Figure 1 D-F). Neutrophils from dSSc patients also exhibited greater spontaneous ROS production and enhanced migration toward chemoattractant (Figure 1 G, H, Supplemental Figure 1 B for representative flow cytometry plots). Biopsy tissue from lesional skin revealed more neutrophils in patients than controls (Supplemental Figure 1C and Supplemental Table 3 for demographics). Thus, in dSSc, neutrophils are abundant in blood, exhibit an activated phenotype in circulation, and can be identified in affected tissue.

**FIGURE 1.**
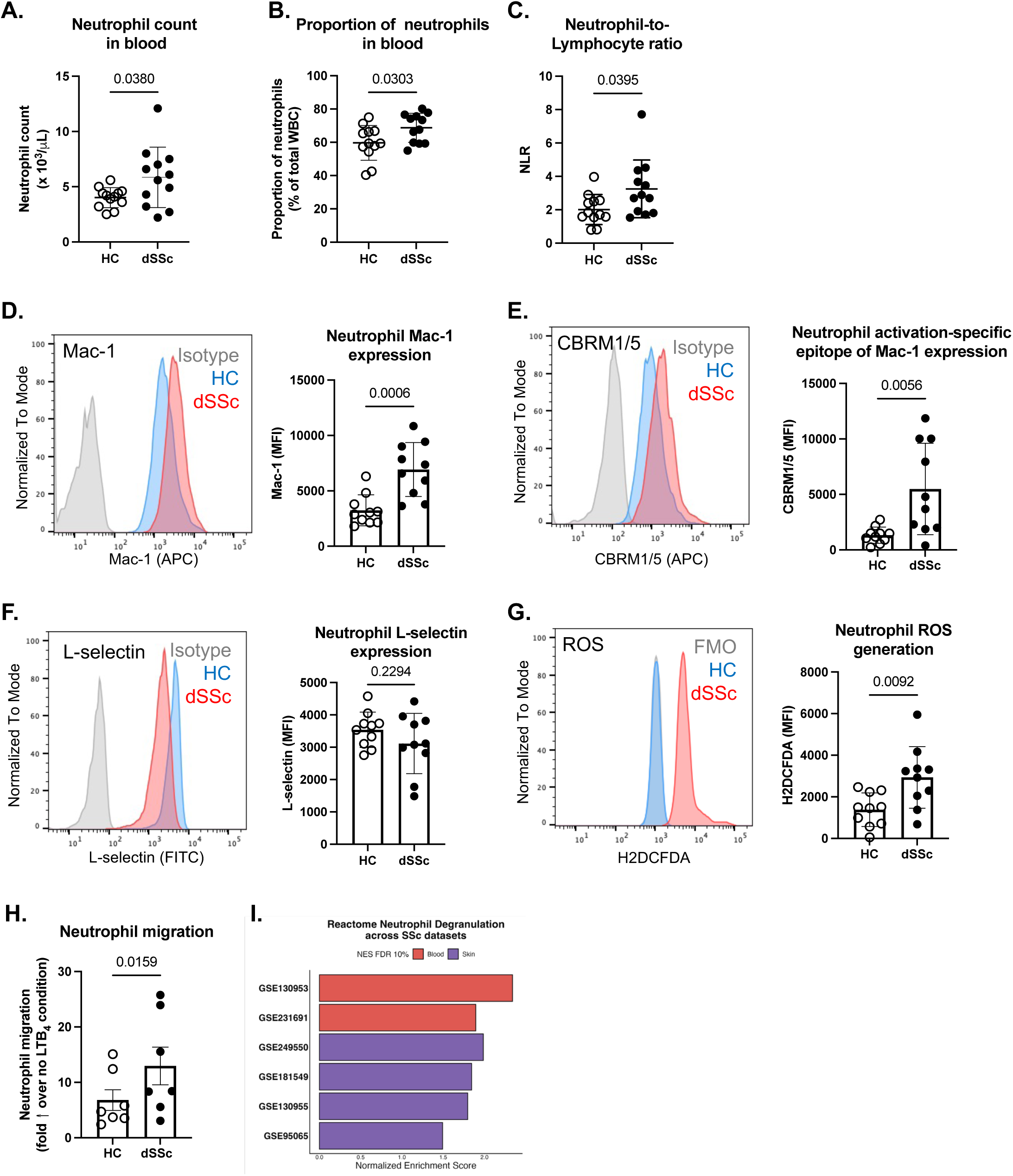
Neutrophils exhibit an activated phenotype in human dSSc. **(A-C)** Complete blood count (CBC) was performed in 12 healthy controls (HC) and 12 patient with diffuse systemic sclerosis (dSSc). **(A)** Neutrophil count (x10^3^/mL) in whole blood of HC and dSSc. **(B)** Proportion of neutrophils among total white blood cells (WBC) in HC and dSSc. **(C)** Neutrophil-to-Lymphocyte ratio in HC and dSSc. **(D-G)** Whole blood neutrophils from dSSc or healthy controls were assayed by flow cytometry. Representative histogram plot (left) and quantification (right) showing expression of the indicated markers in neutrophils from HC and dSSc patients (n=10 per group). **(H)** Neutrophils from HC and dSSc (n= 7/group) were assayed for their migration capacity toward the chemoattractant LTB_4_. Data are expressed as fold increased over no LTB_4_. Each dot is the average of duplicates for each condition. **(I)** Reactome Neutrophil Degranulation Pathway. Normalized Enrichment Score (NES) for the *Neutrophil Degranulation* pathway across all analyzed SSc datasets that met the 10% threshold. Bars are colored by tissue (blood in red, skin in purple). All datasets showed consistent enrichment, with stronger signals observed in blood.

To test the generalizability of these findings, we examined 6 publicly available transcriptomic datasets (2 from blood and 4 from skin biopsy samples) comparing SSc patients (dSSc when specified by the dataset) with controls for whole blood (121 dSSc, 62 controls) and skin biopsy tissue (184 dSSc, 125 controls) (Supplemental Table 4 for transcriptomic dataset details). Gene Set Enrichment Analysis (GSEA) showed that the *Neutrophil Degranulation* gene set was significantly upregulated in SSc patients compared to healthy controls in both blood and skin, meeting a 10% false discovery rate (FDR) threshold in all datasets, with the strongest enrichment signal observed in the blood (Figure 1 I and Supplemental Figure 2). Differential gene expression analysis of neutrophil-associated genes across the 6 transcriptomic datasets confirmed these observations (Supplemental Figure 3). Together with our flow cytometry and histology data, these findings establish neutrophils as a population of interest in dSSc, justifying further exploration of their contribution.

### Murine models of SSc feature neutrophil activation resembling human dSSc

SSc-like fibrosis of skin and lung mediated through endothelial injury can be induced in wild-type C57Bl/6 mice by intradermal injection of the ROS donor hypochlorous acid (HOCl) or by subcutaneous injection of the chemotherapeutic agent bleomycin (BLM) (*31, 32*). We evaluated the profibrotic role of neutrophils in these two systems, comparing HOCl and BLM mice to controls receiving injections of PBS (Supplemental Figure 4 A & B for model design). Only female animals were employed since male mice developed skin necrosis in the HOCl model and could not be studied humanely (not shown).

At harvest, day 42 (HOCl) and day 28 (BLM), mice exhibited increased dermal thickness (Figure 2 A). Lung fibrosis also developed, as quantified by the modified Ashcroft histological score (*33*), lung collagen content (Figure 2 B & C), and α-smooth muscle actin (α-SMA; Figure 2 D). HOCl and BLM-treated animals developed increased serum anti-Scl-70 autoantibody levels, as previously reported in both models as well as in human dSSc (*34–36*) (Figure 2 E). In both models, circulating neutrophils displayed changes comparable to those we had observed in human patients, including higher surface Mac-1, lower surface L-selectin, greater ROS production, and enhanced migration toward chemoattractant (Figure 2 F-K, Supplemental Figure A-E). In BAL fluid, we observed a fourfold increase in neutrophils in BLM-treated mice compared to controls, although not in the slower-acting HOCl model (Supplemental Figure 4 F & G); both models exhibited a clear increase in neutrophils in perfused and disaggregated lung tissues, as a percentage of all cells and by absolute count (Supplemental Figure 4 H-K). These results support HOCl and BLM as appropriate models to study the pathogenic relevance of neutrophils in dSSc.

**FIGURE 2.**
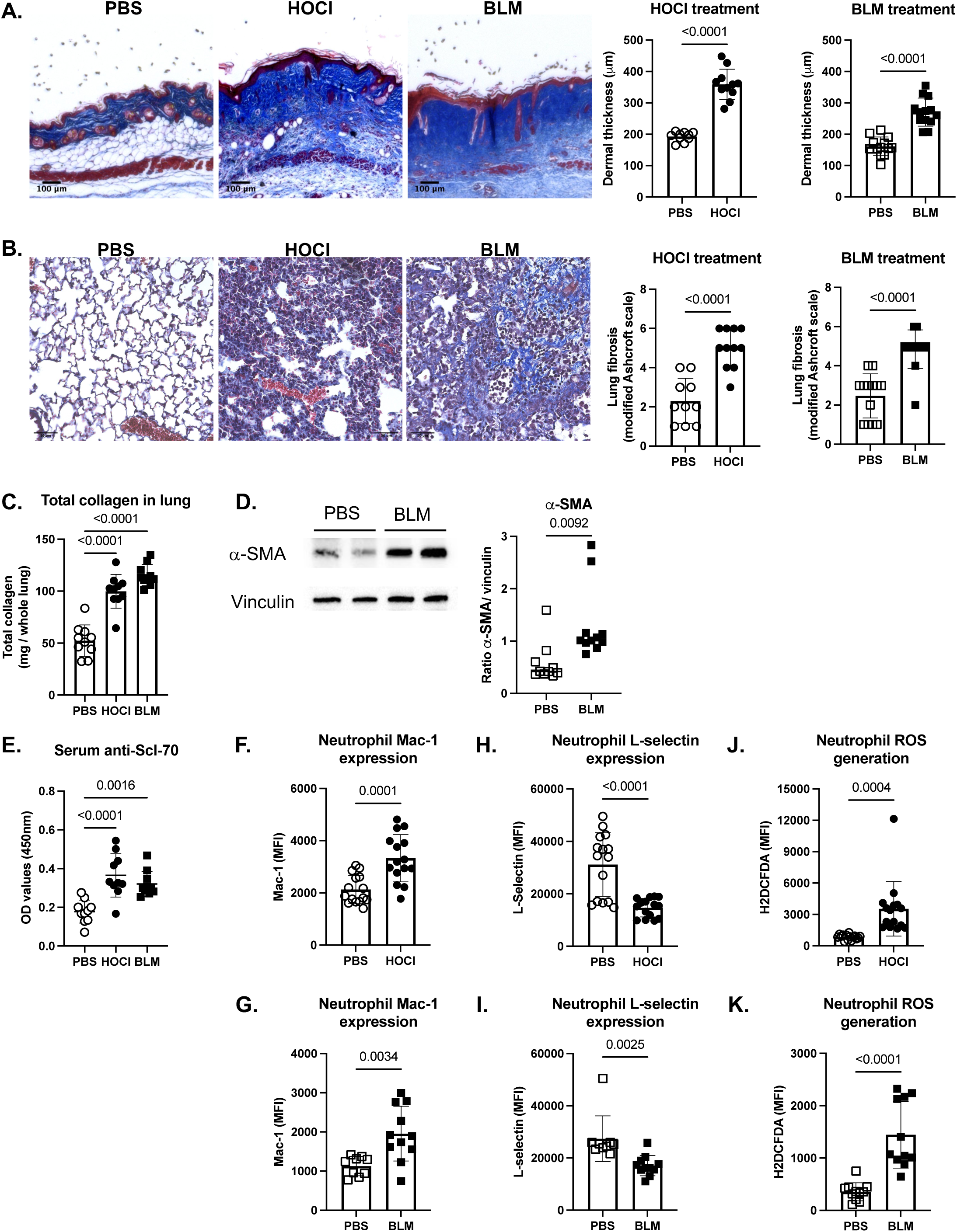
HOCl-SSc and BLM-SSc models recapitulate aspects of human dSSc. **(A)** Representative skin sections (left) of a control skin (PBS) compared to back skin from a d42 and a d28 HOCl- and BLM-SSc mouse (original magnification 4x; Masson Trichrome staining). Dermal thickness (right) was measured for each group (HOCl model: n = 10 PBS and n =11 HOCl; BLM model: n= 13 PBS and n=13 BLM). **(B)** Representative lung sections (left) of a control lung (PBS) compared to lung from d42 and a d28 HOCl- and BLM-SSc mice, respectively (original magnification 20x; Masson Trichrome staining). Microscopic scores of lung sections according to a modified Ashcroft’s method (right) were determined for each group (HOCl model: n = 10 PBS and n =11 HOCl; BLM model: n= 12 PBS and n=13 BLM). **(C)** Total collagen content in the lung from PBS-, HOCl- or BLM-treated mice (n=10 per group). **(D)** Western blot analysis of α-SMA in lung lysates of PBS or BLM-treated mice (representative pictures show on the left and quantification on the right). **(E)** Level of anti-Scl-70 antibodies in the serum of PBS-, HOCl- or BLM-treated mice (n=10 per group). **(F-K)** Whole blood neutrophils from HOCl- or BLM-SSc or PBS control mice were assayed by flow cytometry (HOCl model: n = 15 PBS and n =15 HOCl; BLM model: n= 15 PBS and n=15 BLM). **(F)** Quantification of Mac-1 expression in neutrophils from PBS- or HOCl-treated mice. **(G)** Quantification of Mac-1 expression in neutrophils from PBS- or BLM-treated mice. **(H)** Quantification of L-selectin expression levels in neutrophils from PBS- or HOCl-treated mice. **(I)** Quantification of L-selectin expression levels in neutrophils from PBS- or BLM-treated mice. **(J)** Quantification of ROS levels in neutrophils from PBS- or HOCl-treated mice. **(K)** Quantification of ROS levels in neutrophils from PBS- or BLM-treated mice. The data represent the mean ± SD for all experiments. HOCl, hypochlorous acid; BLM, bleomycin; ROS, reactive oxygen species; FMO, fluorescence minus one.

### Neutrophils mediate fibrosis in murine models of SSc

To test the role of neutrophils in the development and progression of fibrosis in experimental SSc, we depleted neutrophils by intraperitoneal (i.p.) administration of 100 μg of anti-Ly6G antibody (clone 1A8) vs. irrelevant isotype control (clone 2A3) daily for 7 days then every 48 hours. Starting from day 7, mice received HOCl or BLM to induce fibrosis (Supplemental Figure 6 A). Neutrophil depletion (∼80%) was sustained for up to 49 days, markedly attenuating dermal thickness and lung fibrosis in both models (Figure 3 A-G, Supplemental Figure 6 B).

**FIGURE 3.**
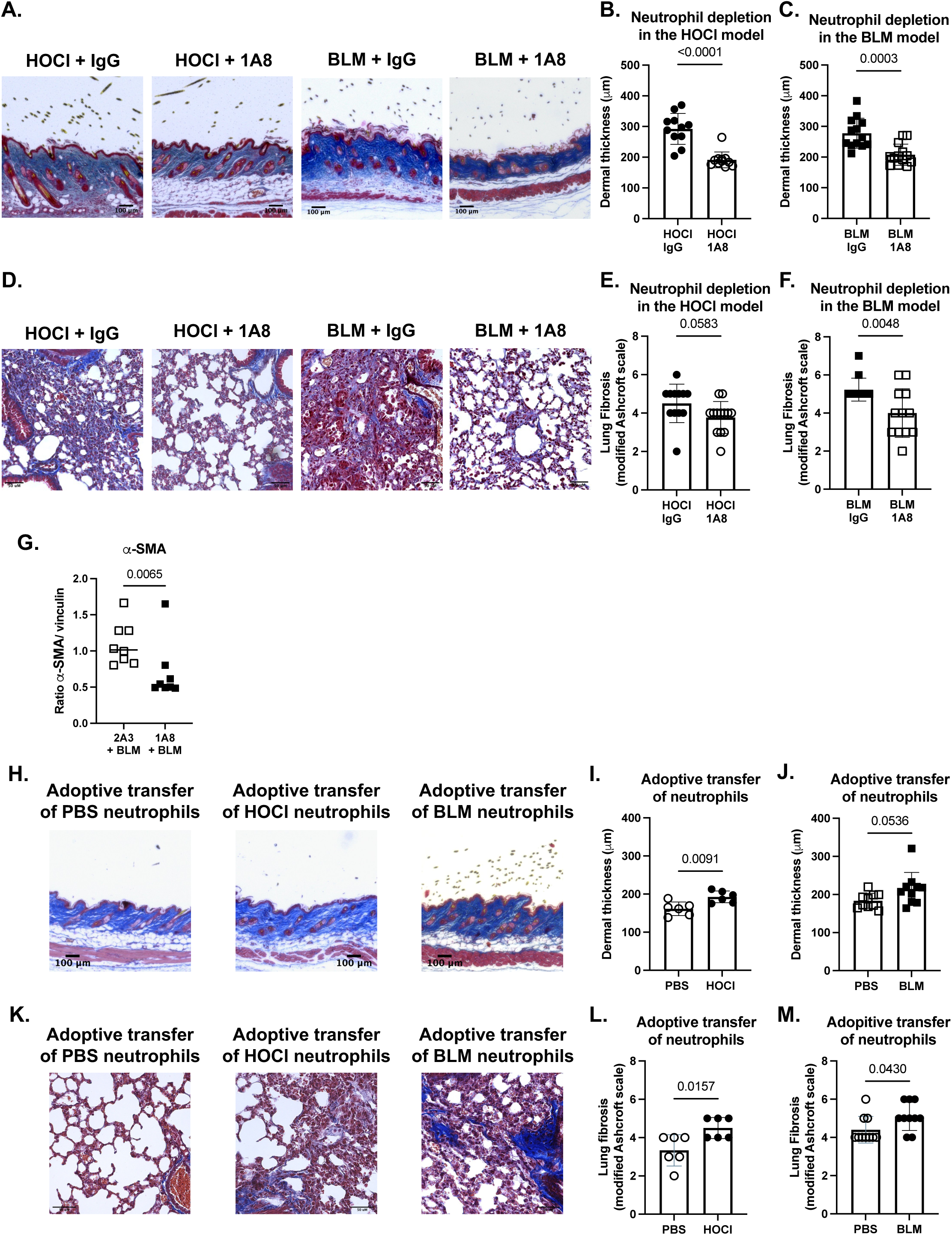
Neutrophils mediate fibrosis in mouse SSc. **(A)** Representative skin sections isolated from HOCl or BLM -mice treated with IgG or with 1A8, respectively (original magnification 4x; Masson Trichrome staining). **(B & C)** Dermal thickness was measured for HOCl mice treated with IgG or 1A8 (n=12 IgG and n= 13 HOCl) and for BLM-SSc mice treated with IgG or 1A8 (n=13 IgG and n=12 BLM). **(D)** Representative lung sections isolated from HOCl or BLM mice treated with IgG or 1A8 (original magnification 20x; Masson Trichrome staining). **(E & F)** Microscopic scores of the lung sections according to a modified Ashcroft’s method were determined for HOCl mice treated with IgG or 1A8 (n=12 IgG and n= 13 HOCl) and for BLM-SSc mice treated with IgG or 1A8 (n=13 IgG and n=12 BLM). **(G)** Quantification of Western blot analysis of α-SMA in lung lysates of BLM-mice treated with IgG or with 1A8**. (H)** Representative skin sections isolated from WT mice 2 weeks following adoptive transfer (i.d injection) of neutrophils isolated from PBS-, HOCl- or BLM-treated mice (original magnification 4x; Masson Trichrome staining). **(I-J)** Dermal thickness was measured for each group (HOCl model: n = 6 PBS and n = 6 HOCl; BLM model: n= 10 PBS and n=10 BLM). **(K)** Representative lung sections isolated from wild-type mice 2 weeks following adoptive transfer (i.v. injection) of neutrophils isolated from PBS-, HOCl- or BLM-treated mice (original magnification 20x; Masson Trichrome staining). **(L-M)** Microscopic scores of the lung sections according to a modified Ashcroft’s method were determined for each group (HOCl model: n = 6 PBS and n = 6 HOCl; BLM model: n= 10 PBS and n=10 BLM). The data represent the mean ± SD.

To test whether neutrophils directly contribute to fibrosis, we employed adoptive transfer. Neutrophils were isolated by negative selection from bone marrow of mice treated with PBS (day 28 or 42), HOCl (day 42), or BLM (day 28); 5×10^6^ cells were then injected into heathy recipients intradermally (i.d.) to test an effect on skin or intravenously (i.v.) to test an effect on lungs, and tissues were harvested after 2 weeks (Supplemental Figure 7 A & B). Remarkably, neutrophils from HOCl- and BLM-treated mice induced fibrosis in the skin and lungs of healthy mice compared with those from PBS-treated animals (Figure 3 H-M). As a methodological control, we transferred isolated T cells from PBS- or HOCl-treated mice; these cells failed to induce fibrosis (Supplemental Figure 7 C-G). Together, these findings identify neutrophils as both necessary and sufficient for induction of fibrosis in experimental SSc.

### Neutrophils release NETs to mediate fibrosis in experimental SSc

Next, we sought to determine how neutrophils mediate tissue fibrosis. Among neutrophil effector functions, formation of NETs is particularly interesting. NETs promote inflammation through mechanisms including induction of pro-inflammatory cytokines and chemokines (*37, 38*), activation of immune cells (*39, 40*), and generation of oxidative stress (*41, 42*). NETs also induce epithelial-mesenchymal transition (*43–45*), activate fibroblasts (*46, 47*), and facilitate extracellular matrix deposition (*48, 49*). Human patients with SSc exhibit elevated circulating NET remnants (*15, 30*). We observed that dSSc neutrophils stimulated with the calcium ionophore A23187 released more NETs than those from control donors, as assessed by basal CitH3 release; interestingly, dSSc but not healthy neutrophils spontaneously generated NETs during culture (Figure 4 A, Supplemental Figure 8 A). We therefore explored whether neutrophils contribute to SSc via NETs.

**FIGURE 4.**
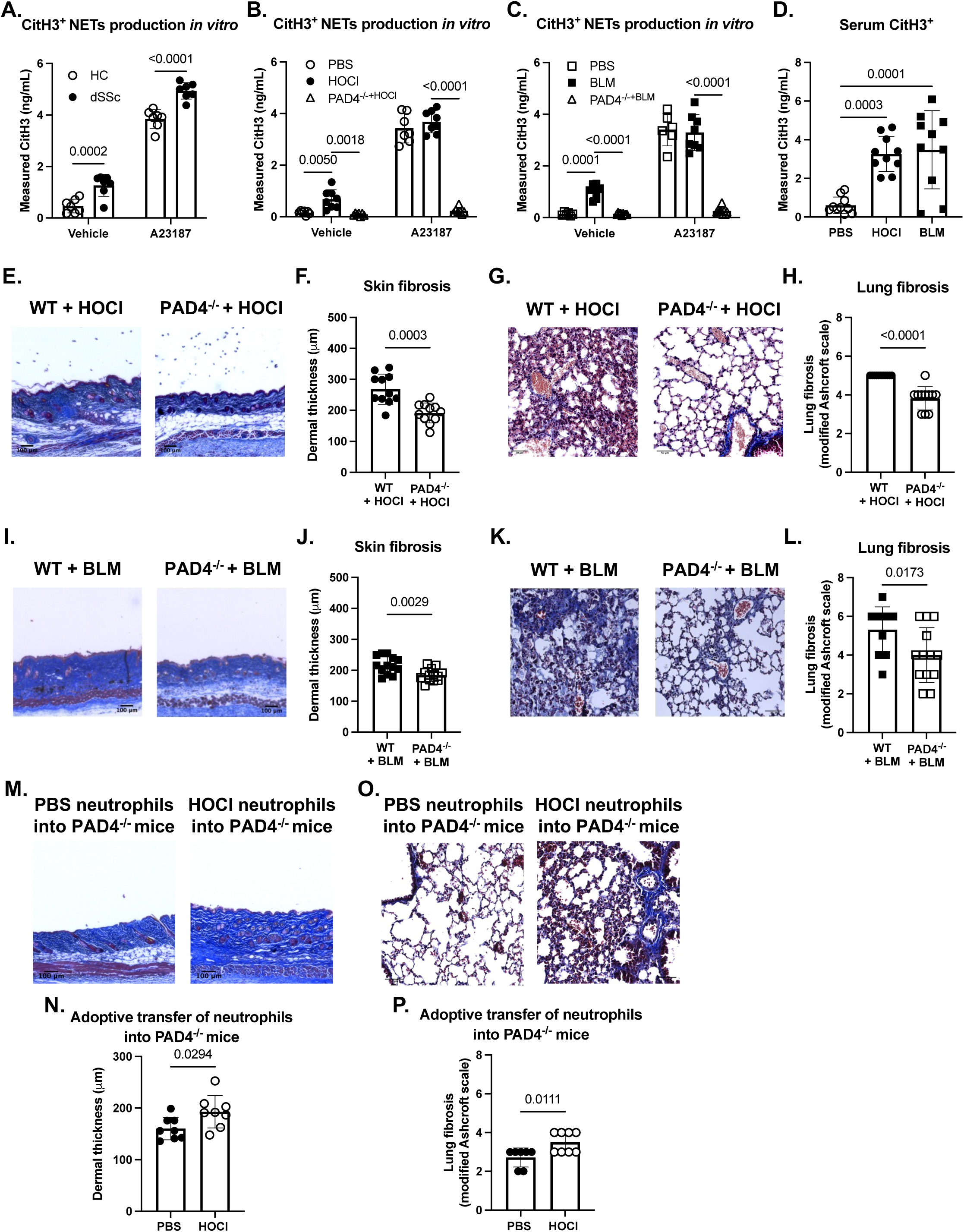
NETs released by neutrophils participate in fibrosis in SSc. **(A)** NET generation by human neutrophils was assayed *in vitro* after treatment with vehicle (DMSO) or A23187 (1μM) for 4h, assessing NET-bound CitH3 by ELISA (n= 7/group); each dot represents an independent experiment. **(B)** Levels of NET-bound CitH3 analyzed by ELISA from neutrophils isolated from wild-type mice treated with PBS (n=7) or HOCl (n=8) or from PAD4^-/-^(n=7) mice treated with HOCl. **(C)** Levels of NET-bound CitH3 analyzed by ELISA from neutrophils isolated from WT mice treated with PBS (n= 6) or BLM (n= 8) or from PAD4^-/-^ (n=9) mice treated with BLM. **(D)** Levels of CitH3 in serum of PBS, HOCl or BLM mice analyzed by ELISA (n=10 per group). **(E)** Representative skin sections isolated from WT or PAD4^-/-^ mice treated with HOCl, respectively (original magnification 4x; Masson Trichrome staining). **(F)** Dermal thickness was measured for each group (n=11 per group). **(G)** Representative lung sections isolated from WT or PAD4^-/-^ mice treated with HOCl (original magnification 20x; Masson Trichrome staining). **(H)** Microscopic scores of lung sections according to a modified Ashcroft’s method (right) were determined for each group (n=11 per group). (**I**) Representative skin sections isolated from WT or PAD4^-/-^ mice treated with BLM, respectively (original magnification 4x; Masson Trichrome staining). (**J**) Dermal thickness was measured for each group (n=13 per group). (**K**) Representative lung sections isolated from WT or PAD4^-/-^ mice treated with BLM (original magnification 20x; Masson Trichrome staining). (**L**) Microscopic scores of lung sections according to a modified Ashcroft’s method (right) were determined for each group (n=13 per group). **(M)** Representative skin sections isolated from PAD4^-/-^ mice 2 weeks following adoptive transfer of neutrophils isolated from PBS- or HOCl-treated mice (original magnification 4x; Masson Trichrome staining). **(N)** Dermal thickness was measured for each group (n=8 per group). **(O)** Representative lung sections isolated from PAD4^-/-^ mice 2 weeks following adoptive transfer of neutrophils isolated from PBS- or HOCl-treated mice (original magnification 20x; Masson Trichrome staining). **(P)** Microscopic scores of the lung sections according to a modified Ashcroft’s method were determined for each group (n=7 PBS and 8 HOCl). The data represent the mean ± SD. CitH3, citrullinated H3; NETs, neutrophil extracellular traps; PAD4, Peptidylarginine deiminase 4; WT, wild type.

In both HOCl- and BLM-treated mice, neutrophils exhibited enhanced spontaneous NET release, although unlike human dSSc no difference was observed after stimulation with the calcium ionophore A23187 (Figure 4 B & C). NET formation requires chromatin decondensation (*50*), mediated through histone H1, H3 and H4 deimination by peptidyl arginine deiminase 4 (PAD4), an enzyme highly expressed in neutrophils (*51*). In all conditions, CitH3 was essentially absent in neutrophils from PAD4^-/-^ animals, confirming the specificity of the assay for NETosis (Figure 4 B & C).

*In vivo*, mice subjected to HOCl and BLM treatment exhibited a notable increase in NETosis, as assessed by the levels of CitH3 in both serum and lung tissue (Figure 4 D and Supplemental Figure 8 B). PAD4^-/-^ mice exhibited markedly reduced skin and lung fibrosis in both HOCl and BLM mice, with levels of CitH3 reduced to control levels in lung homogenates (Figure 4 E-L and Supplemental Figures 8 B). Correspondingly, total collagen content in whole lungs was reduced in PAD4^-/-^ mice compared to WT mice (HOCl mice, Supplemental Figure 8 C). Adoptive transfer of neutrophils from HOCl mice induced skin and lung fibrosis in healthy PAD4^-/-^ recipients, confirming a specific role for neutrophil NETs as opposed to other effects of PAD4 deficiency (Figure 4 M-P). Interestingly, PAD4 deficiency did not reduce neutrophil activation as assessed by surface Mac-1 and L-selectin, although PAD4^-/-^ mice did show reduced basal ROS production (Supplemental Figure 8 D-I). These data implicate NET formation as a critical endpoint of neutrophil activation contributing to fibrosis in murine SSc.

### Platelets are activated during SSc and participate in fibrosis in murine models

We sought to understand how neutrophils become activated during SSc. Our attention focused on platelets because of their established role in neutrophil activation and migration (*52–54*). GSEA showed an enrichment in platelet-related activation pathways across the 6 transcriptomic datasets in SSc, all meeting the 10% FDR threshold, with again a higher NES for platelet pathways in the blood datasets (Figure 5A, Supplemental Figure 9). In fresh citrate-anticoagulated whole blood from patients with dSSc, platelets exhibited increased basal surface P-selectin and phosphatidyl serine expression compared with healthy controls (Figure 5 B & C). Comparable changes were observed in mice treated with HOCl or BLM, showing that activation of circulating platelets is a feature shared across human SSc and murine SSc models (Figure 5 D-G, Supplemental Figure 10 A-C for gating strategy).

**FIGURE 5.**
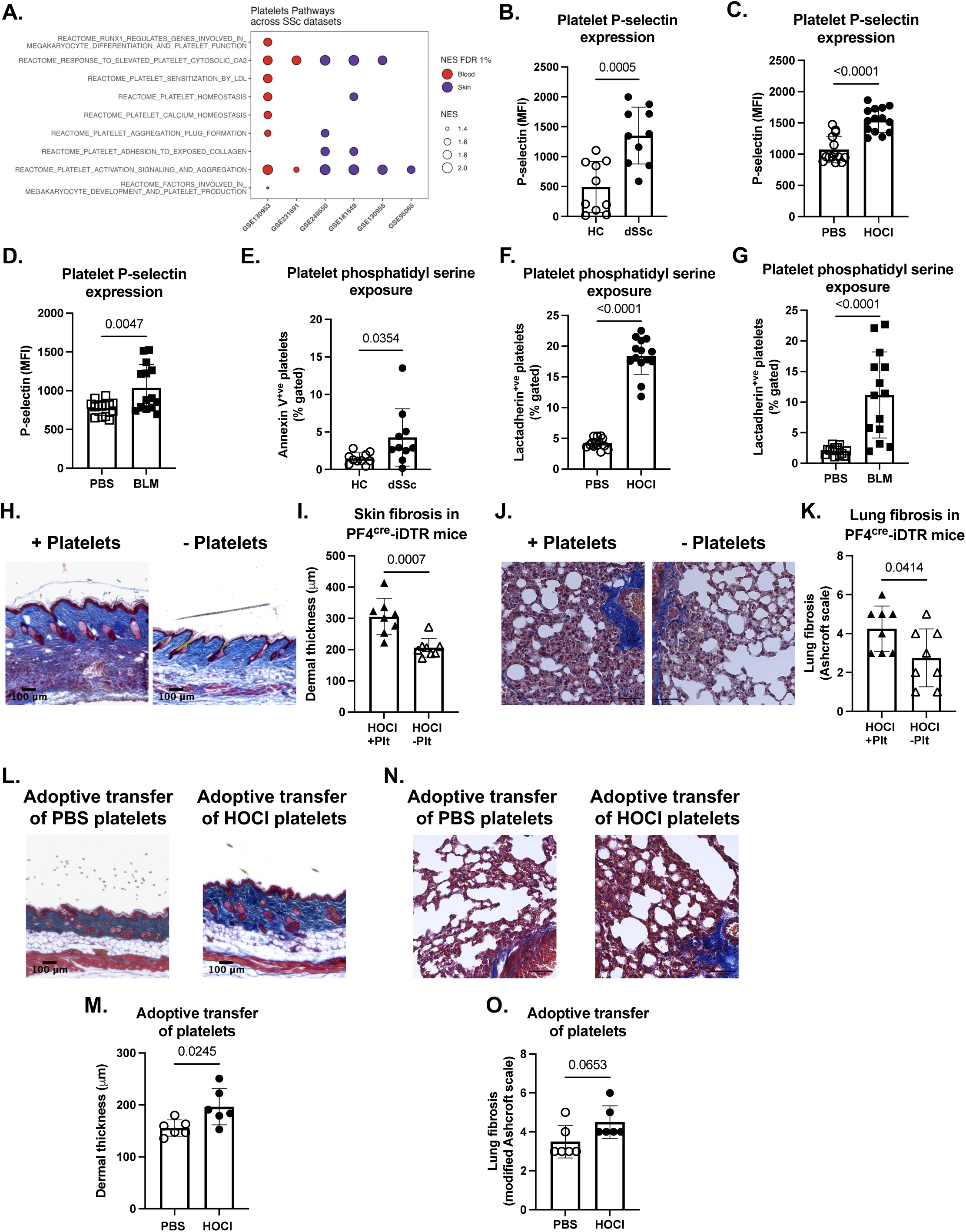
Platelets are activated during SSc and participate in fibrosis. **(A)** Reactome Platelet Pathways. Normalized Enrichment Scores (NES) for platelet-related pathways across transcriptomic datasets in SSc meeting the 10% threshold. Dot color indicates tissue of origin (blood in red, skin in purple), and dot size reflects the magnitude of the NES. **(B-F)** Whole blood platelets from dSSc patients and healthy controls (n=10 per group) and HOCl-SSc or BLM-SSc or PBS control mice (n=14 per group) were assayed by flow cytometry. **(B-D)** Quantification of the P-selectin expression levels on platelets from human dSSc patients **(B)**, mouse HOCl-SSc **(C)** or mouse BLM-SSc **(D)**, compared to healthy controls for each group. **(E-G)** Quantification of phosphatidyl serine expression on platelet surface from human dSSc patients **(E)**, mouse HOCl-SSc **(F)** or mouse BLM-SSc **(G)**, compared to healthy controls for each group. **(H)** Representative skin sections isolated from Pf4^cre-^-iDTR^fl/fl^ (+ platelets) or Pf4^cre+^-iDTR^fl/fl^ (-platelets) mice treated with HOCl (original magnification 4x; Masson Trichrome staining). **(I)** Dermal thickness was measured for each group (n=8 per group). **(J)** Representative lung sections isolated from Pf4^cre-^-iDTR^fl/fl^ (+ Platelets) or Pf4^cre+^-iDTR^fl/fl^ (-Platelets) mice treated with HOCl (original magnification 20x; Masson Trichrome staining). **(K)** Microscopic scores of lung sections according to a modified Ashcroft’s method (right) were determined for each group (n=8 per group). **(L)** Representative skin sections isolated from WT mice 2 weeks following adoptive transfer of platelets isolated from PBS- or HOCl-treated mice (original magnification 4x; Masson Trichrome staining). **(M)** Dermal thickness (was measured for each group (n=6 per group). **(N)** Representative lung isolated from WT mice 2 weeks following adoptive transfer of platelets isolated from PBS- or HOCl-treated mice (original magnification 20x; Masson Trichrome staining). **(O)** Microscopic scores of lung sections according to a modified Ashcroft’s method (right) were determined for each group (n=6 per group). The data represent the mean ± SD.

To test whether platelets play a causal role in SSc, we generated mice in which expression of the diphtheria toxin (DT) receptor is expressed selectively in megakaryocytes and their daughter platelets (PF4^cre^-iDTR), allowing near-total platelet depletion by administration of DT (*55*). We administered DT daily for 7 days then every 72h into PF4^Cre+^/DTR^fl/fl^ mice and PF4^Cre-^/DTR^fl/fl^ littermate controls, initiating the HOCl model on day 7 (Supplemental Figure 10 D for experimental design). Platelet depletion (>99%) was sustained for up to 49 days (Supplemental Figure 10 E). On day 42 following HOCl injections, dermal thickness was markedly decreased in platelet-depleted PF4^Cre+^/DTR^fl/fl^ compared to littermate controls treated with DT (Figures 5 H & I). Similarly, lung fibrosis was markedly reduced in mice subjected to platelet depletion (Figures 5 J & K).

To further test the profibrotic role of platelets, we performed adoptive transfer. Platelets isolated from HOCl- or PBS-treated mice were injected (5×10^6^ platelets) i.d. or i.v. into heathy recipient animals (Supplemental Figure 7 A). After 2 weeks, skin and lung tissues were collected for histopathological evaluation. The transfer of platelets from HOCl-treated mice induced increased dermal thickness and a trend toward increased lung fibrosis compared to mice receiving platelets isolated from control PBS-treated mice (Figure 5 L-M). Collectively, these findings confirm participation of platelets in fibrosis in experimental SSc.

### Platelets trigger the profibrotic function of neutrophils in experimental SSc

To evaluate whether the effect of platelets could be mediated through neutrophils, we assessed the kinetics of platelet and neutrophil activation in the slower HOCl model. Blood was collected at days 0, 14, 28 and/or 42 following initiation of HOCl or PBS treatment. Platelet activation was assessed by surface expression of P-selectin and phosphatidyl serine, while neutrophil activation was monitored by surface expression of Mac-1 and L-selectin as well as intracellular ROS. We observed that platelet activation began between day 14-28, preceding neutrophil activation by at least 2 weeks (Figure 6 A-E). In PF4^Cre+/^DTR^fl/fl^ mice, platelet depletion reduced neutrophil activation, as reflected in decreased Mac-1 levels and increased L-selectin (Figures 6 F & G). These findings position platelet activation prior to, and so potentially contributory to, neutrophil activation.

**FIGURE 6.**
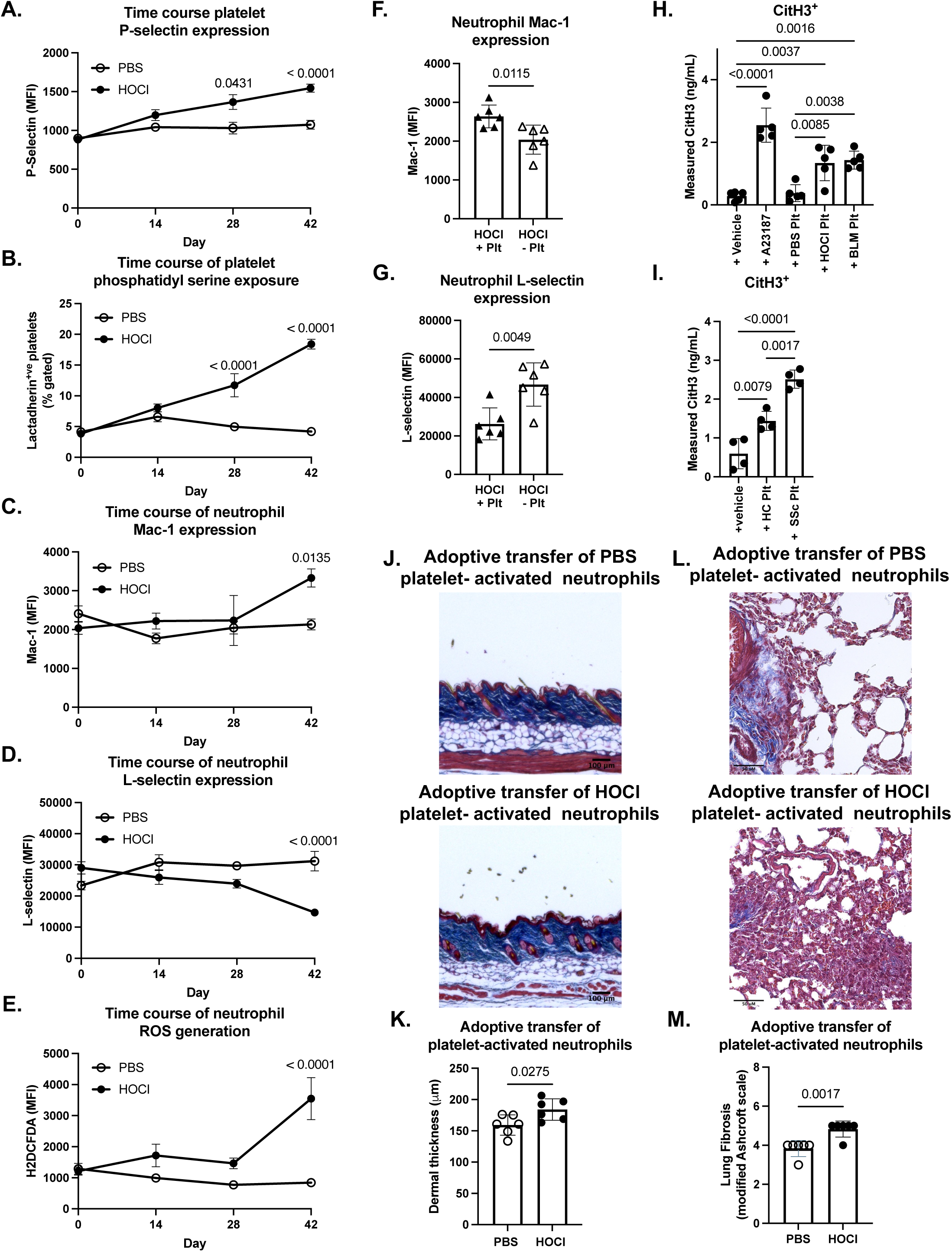
Platelets trigger the profibrotic function of neutrophils in SSc. **(A-E)** Whole blood platelet and neutrophil activation were assayed by flow cytometry (n=15 per group). Kinetics of activation was assessed every 14 days in PBS and HOCl mice and platelet activation was monitored via change in P-selectin **(A)** and phosphatidyl serine **(B)** expression while neutrophil activation was monitored via change in the expression of Mac-1 **(C)** and L-selectin **(D)** and in ROS generation **(E)**. (**F-G**) Mac-1 **(F)** and L-selectin **(G)** expression were assessed on neutrophils from Pf4^cre-^-iDTR^fl/fl^ (+ platelets) or Pf4^cre+^-iDTR^fl/fl^ (-platelets) mice treated with HOCl (n=6 per group). **(H)** Levels of NET-bound CitH3 analyzed by ELISA from WT neutrophils co-cultured for 4h with platelets isolated from PBS-, HOCl- or BLM-treated mice. Vehicle (DMSO) or A23187 (1μM) serve as control (n=5 per group). **(I)** Levels of NET-bound CitH3 analyzed by ELISA from healthy naive neutrophils co-cultured for 3h with platelets isolated from healthy control or dSSc patients. Vehicle (untreated) serves as control (n=4 per group). **(J)** Representative skin sections isolated from WT mice 2 weeks following adoptive transfer of neutrophils pre-incubated with platelets isolated from PBS or HOCl-treated mice (original magnification 4x; Masson Trichrome staining). **(K)** Dermal thickness was measured for each group (n=6 per group). **(L)** Representative lung sections isolated from WT mice 2 weeks following adoptive transfer of neutrophils pre-incubated with platelets isolated from PBS- or HOCl-treated mice (original magnification 20x; Masson Trichrome staining). **(M)** Microscopic scores of lung sections according to a modified Ashcroft’s method (right) were determined for each group (n=6 per group). The data represent the mean ± SD. Plt, platelet.

To test whether platelets act directly on neutrophils to enable SSc, we assessed the ability of platelets from mice treated with PBS, HOCl, or BLM to induce NETs in healthy neutrophils. Neutrophils isolated from healthy mice were co-cultured with HOCl, BLM or PBS control platelets (1 neutrophil: 50 platelets). Vehicle and A23187 were used as negative and positive control, respectively. Indeed, platelets from HOCl and BLM mice induced NET formation as measured by released CitH3 (Figure 6 H). This finding was recapitulated with human samples, demonstrating that platelets from dSSc donors trigger NET formation in healthy neutrophils more than platelets from healthy donors (Figure 6 I).

These findings suggest that platelet-activated neutrophils could potentially mediate disease *in vivo*. To confirm this possibility, we performed adoptive transfer experiments using healthy neutrophils pre-incubated with HOCl platelets and then washed. Indeed, compared with neutrophils incubated with control platelets, neutrophils exposed to HOCl platelets increased dermal thickness and lung fibrosis in previously healthy mice (Figure 6 J-M). Thus, activated platelets from murine SSc engage neutrophils as profibrotic effector cells via mechanisms that may include NETs.

### Feedback activation of platelets by neutrophils in experimental SSc

Neutrophils can promote platelet activation (*56, 57*). Correspondingly, HOCl and BLM mice depleted of neutrophils by 1A8 exhibited reduced platelet activation compared to those treated with 2A3, as reflected in lower expression of P-selectin and phosphatidyl serine (Supplemental Figures 11 A-D). PAD4^-/-^ mice treated with HOCl or BLM similarly exhibited reduced platelet activation compared with controls (Supplemental Figure 11 E-H), suggesting that feedback from neutrophils to platelets is mediated via NETs, though reduced platelet activation might also result from other processes, such as attenuated tissue injury.

### The platelet-neutrophil axis of fibrosis in murine SSc is initiated via the GPVI receptor

We sought to determine how platelets become activated in SSc. Microvascular injury is an early disease feature, leading to exposure of collagen beneath the damaged endothelium (*58–60*). Some but not all authors have identified a specific enhanced reactivity of human SSc platelets to collagen (*27–29*). In our differential expression analysis of platelet-associated genes across the 6 transcriptomic datasets in SSc, several genes involved in the collagen receptor glycoprotein VI (GPVI) pathway emerged as significantly enriched, including LYN, SYK and PLCg2 (Supplemental Figure 9). We isolated platelets from 3 dSSc patients and sex-matched healthy donors and tested reactivity to ADP (agonist for the P2Y1 and P2Y12 receptors), thrombin (agonist for the protease activation receptors), and convulxin (agonist for GPVI). We observed a trend toward increased reactivity to convulxin, though this effect was obscured by substantial donor-to-donor variation (Supplemental Figures 12 A & B). In HOCl mice, selective platelet reactivity to GPVI ligation emerged clearly (Supplemental Figures 12 C & D). We therefore sought to test the role of GPVI in experimental SSc.

To this end, we subjected GPVI^-/-^ and littermate control GPVI^+/+^ mice to HOCl- and BLM-induced SSc. Flow cytometry confirmed GPVI deficiency in platelets of GPVI^-/-^ mice (Supplemental Figure 12 E & F). GPVI^-/-^ platelets exhibited reduced P-selectin and phosphatidyl serine expression (Figure 7 A & B, Supplemental Figure 13 A & B), consistent with impaired platelet activation. We observed a concurrent reduction in neutrophil activation, with lower Mac-1 expression and higher L-selectin expression (Figure 7 C & D, Supplemental Figure 13 C & D). Serum CitH3 levels were also lower, indicating reduced NET formation (Supplemental Figure 13 E). Histopathological examination of skin and lung sections confirmed a protective effect against fibrosis in GPVI^-/-^ mice in both HOCl and BLM models (Figure 7 E-H).

**FIGURE 7.**
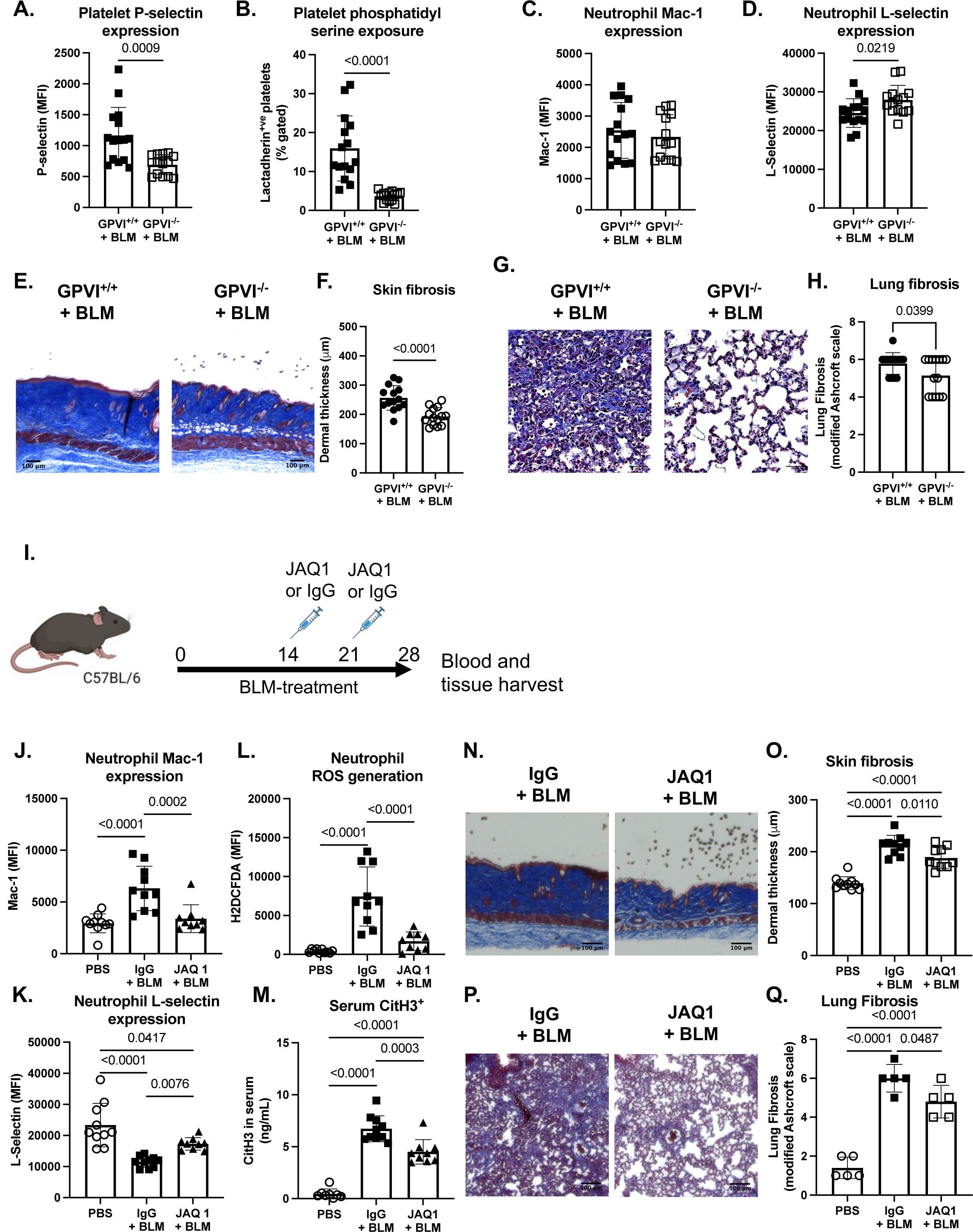
The platelet-neutrophil axis of fibrosis in SSc is activated via the GPVI receptor. **(A-D)** Whole blood platelets and neutrophils were assayed by flow cytometry in GPVI^-/-^ or littermate control GPVI^+/+^ mice treated with BLM at d28 (n= 15 PBS and n=14 BLM). Platelets were assayed for P-selectin and phosphatidyl serine expression in the BLM model **(A & B)**. Neutrophils were assayed for Mac-1 and L-selectin expression in the BLM model **(C & D). (E)** Representative skin sections isolated from GPVI^+/+^ or GPVI^-/-^ mice treated with BLM (original magnification 4x; Masson Trichrome staining). **(F)** Dermal thickness was measured for each group (n=14 per group). **(G)** Representative lung sections isolated from GPVI^+/+^ or GPVI^-/-^ mice treated with BLM (original magnification 20x; Masson Trichrome staining). **(H)** Microscopic scores of lung sections according to a modified Ashcroft’s method (right) were determined for each group (n=14 per group). **(I)** Timeline representing the injection days of JAQ1 or isotype control IgG for GPVI inhibition in the BLM model. (**J-L**) Whole blood neutrophils were assayed by flow cytometry in BLM mice treated with JAQ1 or IgG at d28. Neutrophils were assayed for Mac-1 (**J**) and L-selectin expression (**K**) and ROS generation (**L**) (PBS: n=10, IgG: n= 10, JAQ1: n=9). (**M**) Levels of CitH3 in serum of BLM mice treated with JAQ1 or IgG at d28 was analyzed by ELISA (PBS: n=8, IgG: n= 10, JAQ1: n=9). (N) Representative skin sections isolated from BLM mice treated with JAQ1 or IgG at d28 (original magnification 4x; Masson Trichrome staining). **(O)** Dermal thickness was measured for each group (n=10 per group). **(P)** Representative lung sections isolated from BLM mice treated with JAQ1 or IgG at d28 (original magnification 20x; Masson Trichrome staining). **(Q)** Microscopic scores of the lung sections according to a modified Ashcroft’s method (right) were determined for each group (n=10 per group). The data represent the mean ± SD.

Finally, we examined the potential of targeting GPVI as treatment option in experimental SSc. Using the faster BLM model, we induced disease in WT mice, followed by treatment with anti-GPVI mAb JAQ1 or irrelevant control IgG (100μg/ mouse) i.v. on days 14 and 21 (Figure 7 I). JAQ1 antibody inhibits signaling by inducing shedding of GPVI from the platelet surface, and indeed the receptor was essentially absent from platelets in treated mice (Supplemental Figure 13 J). Compared to IgG control mice, JAQ1-treated animals demonstrated reduced platelet and neutrophil activation (Supplemental Figure 13 K & L, Figure 7 J-L) as well as reduced markers of NETosis in serum (Figure 7 M). Skin and lung examination confirmed the protective effect against fibrosis in JAQ1-treated mice compared to controls (Figure 7 N-Q, Supplemental Figure 13 M). These findings confirm that GPVI drives platelet activation, neutrophil activation, NET release, and tissue fibrosis in experimental SSc.

## DISCUSSION

This study defines a GPVI-platelet-neutrophil-NET-fibrosis axis in experimental SSc and provides evidence that this pathway is engaged in human dSSc. Our findings support a model whereby exposed collagen, potentially from injured endothelium (*17, 30*), activates platelets via the GPVI receptor; in turn, these cells activate neutrophils to release profibrotic NETs (Figure 8). Defined mechanistically in two independent murine models that share biologic features with human dSSc, these findings are further corroborated by studies in fresh samples from patients with dSSc and by transcriptomic data from skin and whole blood samples. Together, the findings help to define the processes linking endothelial injury to endo-organ tissue fibrosis.

**FIGURE 8.**
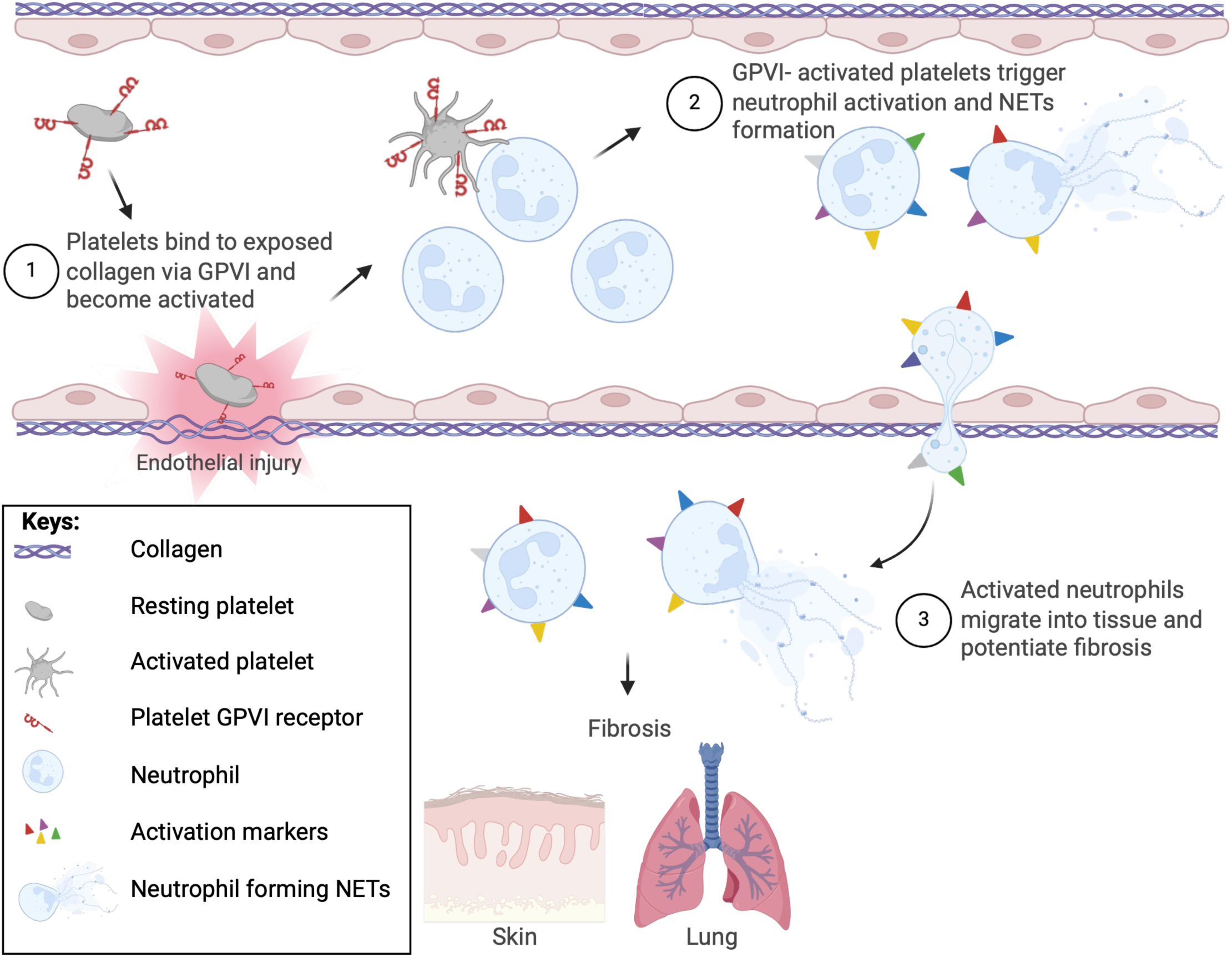
Platelets activate neutrophils to mediate fibrosis in scleroderma. (1) Vascular injury exposes collagen, resulting in activation of circulating platelets via GPVI. (2) Platelets activate neutrophils, with effects including an increased propensity to release neutrophil extracellular traps (NETs). (3) Activated neutrophils and their NETs stimulate fibroblasts to produce fibrosis in skin and lung.

Over the past several decades, multiple groups have observed neutrophil activation in SSc, though some results diverge (*14–18*). This divergence may reflect admixture of dSSc with limited SSc, which is generally less widespread and severe, including fewer circulating endothelial cells as a potential marker of endothelial injury (*61, 62*). Platelet activation in SSc is consistently identified, but it has remained unclear whether this phenotype is incidental or reflective of a pathogenic mechanism, especially since both neutrophils and platelets are rare in SSc tissue biopsies (*17, 28–30*). Using experimental models, we establish platelet and neutrophil activation as potentially critical, linked steps in the pathogenic chain that connects endovascular injury to tissue fibrosis.

Our data show that mice deficient in PAD4 are protected against fibrosis in experimental SSc, aligning with prior research in SSc (*63*). PAD4-mediated histone citrullination enables chromatin decondensation and release (*64*), a process observed predominantly in neutrophils (NETs) but potentially occurring also in cells such as macrophages that may also have a role in SSc-related fibrosis (*65–68*). Using adoptive transfer, we showed that neutrophils are the key source of extracellular traps in fibrosis in our system. These findings do not exclude other effects of neutrophil PAD4 or a role for extracellular traps from other sources. Recently, PAD4 inhibitors have gained interest in the treatment of cancer, rheumatoid arthritis, and COVID (*69–72*). Our findings support the addition of SSc to the list of diseases that may respond to NET blockade.

The observation that the collagen receptor GPVI is central to platelet activation in SSc, and thereby strongly contributory to tissue fibrosis, is particularly interesting. SSc is believed to begin with injury to the endothelium, and our results now support this understanding by showing that a platelet receptor for a component of exposed subendothelial matrix is a critical step leading to neutrophil activation and the elaboration of pro-fibrotic NETs in experimental SSc. This finding is of direct clinical relevance because GPVI is therapeutically targetable in humans. Individuals genetically deficient in GPVI exhibit only a mild bleeding defect and many are asymptomatic, suggesting that interrupting this pathway may be relatively safe (*73, 74*). Some patients with idiopathic thrombocytopenic purpura (*75, 76*) and systemic lupus erythematosus (*77*) exhibit autoantibodies against GPVI, but the resulting clinical phenotype is modest, limited to a mild bleeding tendency and lack of collagen-induced platelet aggregation. Correspondingly, antibodies blocking GPVI are in active clinical development, with preliminary evidence for efficacy and safety in stroke and carotid stenosis (*78–80*). Our findings pave the way for trials of GPVI inhibition in SSc patients, especially since GPVI blockade is not immunosuppressive and therefore might be safely combined with conventional immunomodulatory and anti-fibrotic agents.

The mechanisms by which platelets activate neutrophils are likely complex. Maugeri and colleagues showed that platelet microparticles from SSc patients, and from healthy platelets stimulated with collagen, contain the alarmin HMGB1. This molecule acts directly upon neutrophils to promote autophagy, to persist longer in circulation, and to release NETs. Correspondingly, healthy mice treated with platelet microparticles exhibited evidence of microvasculopathy in lungs, although whether fibrosis follows is undefined (*30*). Our data are fully compatible with these results, showing in two murine models of endothelial-injury-mediated SSc that platelet activation precedes neutrophil activation and that depletion of either neutrophils or platelets prevent disease development. These results confirm that both cell types play key roles in disease development. Activated platelets also stimulate neutrophils through platelet surface P-selectin engaging with its neutrophil ligand PSGL-1, among other mechanisms (*81*). Correspondingly, we observed increased expression of P-selectin in circulating platelets from dSSc patients compared to healthy controls, a finding reflected also in both murine models. Interestingly, P-selectin promotes NET formation (*81*). Other platelet effector functions may also be relevant, both for neutrophil activation and for fibrosis generally. For example, studies in Tsk-1 mice have identified a profibrotic role for platelet serotonin, an effect that presumably proceeds independent of neutrophils (*82*).

We establish that NETs are not only a marker of neutrophil activation in experimental SSc but play a direct pathogenic role. NETs promote fibrosis through various mechanisms, including direct activation of fibroblasts and induction of epithelial-mesenchymal transition, as recently reviewed (*83*). Our data do not address which of these pathways is relevant for SSc; indeed, multiple mechanisms may be involved, and these may vary among tissue sites. Further investigation will be required to identify responsible pathways.

Interestingly, patients with SSc often exhibit elevated levels of D-dimer, resulting from fibrinogen crosslinking and subsequent fibrinolysis, and are at higher risk of venous thrombosis and pulmonary embolism (*84*). The contribution of platelet activation and NET production to this thrombosis risk is unknown. Experimental data confirm that platelet GPVI plays a role in both venous and arterial thrombosis (*85*). NETs also facilitate thrombosis (*86*). These observations suggest that targeting GPVI could potentially have important downstream benefits not only for disease progression but also for thromboembolic risk. The utility of anticoagulation in SSc remains under investigation (*87*).

Our study has several limitations. No murine model perfectly mimics human disease, and extrapolation of findings from the HOCl and BLM models to human SSc must be undertaken with caution. Indeed, human SSc is diverse, in severity, organ involvement, and autoantibody profile (*88*). For this study, we studied patients with dSSc, of whom most had interstitial lung disease, but were not powered to detect differences between groups of patients defined by clinical features or autoantibodies. It is therefore possible that our results are relevant more for some patients than others, and potentially for some organ systems more than others. Further, our dSSc patients differed from controls in demographic features and in ongoing treatment, raising the possibility that some of the differences established here reflected these factors rather than the disease state itself. In this context, it is reassuring that corresponding human neutrophil and platelet phenotypes could be detected in public transcriptional datasets, and that essentially all findings in dSSc neutrophils and platelets were also identified in both HOCl and BLM models. Importantly, endothelial injury – the likely trigger for platelet activation via GPVI through exposure of subendothelial collagen – is a shared feature of all forms of SSc (*89*), irrespective of the upstream autoimmune process inducing the injury. Nevertheless, it will be important to extend these studies by defining platelet and neutrophil phenotypes, as well as circulating NET products, in a broad range of human SSc phenotypes as well as within individual patients before and after treatment.

In conclusion, our study delineates a GVPI-platelet-neutrophil-NETs-fibrosis axis in experimental SSc, together with concordant changes in human dSSC, providing new insights into disease mechanisms and identifying potential new therapeutic targets. Future research will focus on refining these findings and translating them into clinical applications aimed at improving outcomes for SSc patients.

## MATERIAL AND METHODS

### Sex as a biological variable

Human subjects included both males and females. Murine SSc models were conducted exclusively in female mice because male mice develop skin necrosis and cannot be studied humanely. It is unknown whether the findings are relevant for male mice. *In vitro* murine studies included samples from both males and females.

### Animals

C57BL/6J (Jax # 000664), B6.Cg-Padi4tm1.1Kmow/J (PAD4^-/-^, JAX # 030315), C57BL/6-Tg(PF4-icre)Q3Rsko/J (PF4-Cre, JAX # 008535), and C57BL/6-Gt(ROSA)26Sortm1(HBEGF)Awai/J (iDTR, JAX # 007900) mice were purchased from The Jackson Laboratory. Gp6tm1Ware/J (GPVI^-/-^) mice were provided by Professor Jerry Ware. PF4^Cre^-iDTR mice were generated by crossing PF4-icre mice with iDTR mice. The offspring PF4^Cre+^/DTR^fl/fl^ and PF4^Cre-^/DTR^fl/fl^ were used as experimental or littermate control mice, respectively. GPVI^-/-^ mice were bred with C57BL/6 to generate heterozygous GPVI^+/-^ mice. The GPVI^+/-^ mice were them used to generate experimental GPVI^-/-^or GPVI^+/+^ littermate control mice. Genotypes were confirmed using Transnetyx. Both males and females 6-10 weeks old were used for breeding and 6-8-week-old females were used for SSc. Mice were housed under specific pathogen-free conditions within ventilated cages with food and water ad libitum, under constant room temperature and with 12h day-night cycles, at Boston Children’s Hospital. All the procedures were approved under the Institutional Animal Care and Use Committee and operated under the supervision of the department of Animal Resources at Children’s Hospital.

### HOCl-SSc model

HOCl-SSc was induced as previously described (*90*) with slight modification. Briefly, 150 µl of an HOCl solution was intradermally injected in 2 sites (300 µl in total) into the shaved backs of mice, using a 27-gauge insulin syringe 5-days a week for 6 weeks. Control mice received injections of 300 µl of sterile phosphate-buffered saline (PBS). Mice were randomly allocated to experimental and control groups. HOCl was prepared fresh daily by adding NaClO solution (9-14% as active chlorine) to a 100 mM KH_2_PO_4_ solution (pH 6.2). The stock NaClO solution was kept refrigerated and no more than 6 weeks after opening. The NaClO concentration was determined at 292nm using a nanodrop (NanoDrop™ One, Thermo Fisher Scientific) and adjusted between OD 0.7 and 0.9.

### Bleomycin-SSc model

BLM-SSc was induced as previously described (*32*). A sterile solution of 1mg/mL of bleomycin (Cayman chemicals) was in PBS. Bleomycin 100 µl was subcutaneously injected into the shaved backs of mice daily for 4 weeks using a 27-gauge insulin syringe. Control mice received injections of 100 µl of sterile PBS. Mice were randomly allocated into experimental and control groups.

### Tissue collection and histopathologic analysis

Human skin biopsies were collected from the forearm of consented scleroderma or matched healthy control patients. Tissues samples were fixed in 4% PFA for 24h then conserved on 70% ethanol before processing. Sections were stained with an anti-MPO antibody. Slides for histologic assessment were scanned using a EVOS M7000 microscope (Thermo Fisher Scientific) at 4X magnification. The number of neutrophils was assessed by counting the average number of MPO+ neutrophils from 5 randomly selected fields of view (FOV). Assessment was performed independently by 2 blinded experimenters; data shown reflect the mean of both values.

Mouse skin biopsies were collected from the dorsum of the animals at the injection site. Mouse lungs were inflated before collection via insertion of an intratracheal canula with 4% PFA at a flow rate of 200 µL/second (*91*). Tissue samples were fixed in 4% PFA for 24h then conserved on 70% ethanol before processing. Sections (5 mm thick) of paraffin-embedded samples were stained with Masson’s trichrome. Slides for histologic assessment were scanned using a EVOS M7000 microscope (Thermo Fisher Scientific) at 4X magnification. Dermal thickness (layer from the dermal-epidermal junction to the dermal-adipose junction) was calculated from the average of 3 random measurements per section. For mouse lung fibrosis evaluation, average of multiple FOV of the entire lung were scored according to the modified Ashcroft scale defined by Hübner *et al*. (*33*) The scoring criteria were as follows: (A) Grade 0, normal lung. (B) Grade 1, alveoli partly enlarged and rarefied, absence of fibrotic masses, septum thicker than normal. (C) Grade 2, alveoli partly enlarged and rarefied, absence of fibrotic masses, septum thicker than normal with isolated knot-like formation. (D) Grade 3, alveoli partly enlarged and rarefied, absence of fibrotic masses, septum thicker than normal with continuous fibrotic wall in the entire FOV. (E) Grade 4, single fibrotic masses (≤10% of FOV). (F) Grade 5, single fibrotic masses (10-50% of FOV). (G) Grade 6, large contiguous fibrotic masses (>50% of FOV). (H) Grade 7, alveoli nearly obliterated with fibrous masses but some air bubbles. (I) Grade 8, complete obliteration with fibrotic masses. Mouse skin and lung fibrosis assessment was performed blinded to experimenter using ImageJ software (National Institutes of Health).

### Blood collection

Human blood samples were drawn from antecubital veins after providing informed consent, using a 21g butterfly vacutainer tube anticoagulated with citrate or lithium heparin for platelet or neutrophil assessment, respectively. Mouse blood samples were collected as described previously (*92*). Blood was collected from the inferior vena cava of anesthetized animals (isoflurane 1-4%) using a 29g needle attached to a 3mL syringe containing a combination of low molecular weight heparin (enoxaparin, 40 units/ml) and acid-citrate-dextrose (ACD) anticoagulant (13 mM sodium citrate, 1 mM citric acid, 20 mM dextrose, and 10 mM theophylline). For sub-mandibular blood collection, samples were directly collected into heparin tubes.

### Neutrophil enumeration in whole blood

Human neutrophil count in heparin-anticoagulated whole blood was assessed using Sysmex XP-300-Hematology-Analyzer.

### Human neutrophil isolation

Human neutrophils from blood of dSSc patients or sex-matched healthy donors were purified with Ficoll-Paque PLUS (Cytiva). Blood was first diluted with PBS (1:1 ratio) and gently layered on top of 15mL of Ficoll-Paque PLUS onto a 50mL falcon tube. The tube was then centrifuged at 500g for 30 minutes at 18°C-22°C without break. The top 3 layers including plasma, PBMC and Ficoll-Paque PLUS, were removed and discarded. The neutrophil layer directly on top of the red blood cells (RBC) and the first portion of the RBC (approximately 1ml) were collected and placed into a clean 50mL falcon tube. RBC were lysed with ACK (GIBCO, 1:20 blood: ACK ratio) for no more than 2 minutes with constant gentle agitation. A solution of PBS containing 2mM EDTA was added to the tube (1:1 ratio) before centrifugation at 500g for 10 minutes at 18°C-22°C. Supernatant was gently removed and neutrophil pellet resuspended in RPMI 1640 (GIBCO). Neutrophil purity was assessed using Sysmex XP-300 automated hematology analyzer.

### Mouse neutrophil isolation

Bone marrow murine neutrophils were purified using EasySep™ Mouse Neutrophil Enrichment Kit (STEMCELL technologies) according to the manufacturer’s instructions. The purity of neutrophil (Ly6G^+^ subsets) was assessed by flow cytometry.

### T cell isolation

Spleen murine T cells were purified using EasySep™ Mouse T Cell Isolation Kit (STEMCELL technologies) according to the manufacturer’s instructions. The purity of T cells, defined as CD3^+^/CD19^-^, was assessed by flow cytometry.

### Flow cytometry of neutrophil activation

Human heparin-anticoagulated whole blood was diluted with HBSS (1 volume blood: 4 volume HBSS ratio). Diluted whole blood (100μL) was stained at RT for 30 minutes with antibodies and dyes listed in Supplementary Table 2. Thereafter, RBC were lysed with ACK lysis buffer (1 volume blood: 10 volume ACK) for 1 minutes with gentle agitation, centrifugated at 500g for 5 minutes at 4°C and the pellet resuspended in HBSS. Cell suspension was kept on ice until flow cytometry analysis. We characterized human neutrophil activation by assessing levels of Mac-1^+^, L-selectin^+^, activated Mac-1^+^, and ROS (CM-H2DCFDA^+^) on the neutrophil gated CD66b^+^ population. Mouse enoxaparin/ACD-anticoagulated whole blood was diluted with HBSS (1 volume blood: 2 volume HBSS ratio). Mouse samples were treated similarly as human samples with specific antibodies and dyes listed in Supplementary Table 2. We characterized mouse neutrophil activation by assessing levels of Mac-1^+^, L-selectin^+^, and ROS (CM-H2DCFDA^+^) on the neutrophil gated Ly6G^+^ population. For kinetics of activation in HOCl-SSc mice, whole blood was collected via sub-mandibular vein. For each marker, MFI was obtained by background subtraction obtained from isotype control antibody or unstained samples. All samples were acquired with a BD LSRFortessa™ Cell Analyzer (BD biosciences and recorded data analyzed with FlowJo (v.10.10.0).

### Neutrophil migration assay

A volume of 100µL of a suspension of freshly isolated neutrophils (10^6^/mL) in RPMI 1640 medium containing 0.1% FBS was placed in the upper compartment of a transwell chamber featuring uncoated polyester membrane with 3 µm pores (Corning), and a volume of 600 µL of RPMI 1640 medium containing 0.1% FBS and LTB4 (100 nM) or vehicle was added to the bottom compartment. After 2 h incubation at 37°C and 5% CO_2_, 400 µL were harvested from the bottom compartment and neutrophils, determined as the CD66b^+^ (human) or Ly6G^+^ (mouse) population were enumerated by flow cytometry using counting beads.

### Flow cytometry of platelet activation in whole blood

Human heparin-anticoagulated whole blood was diluted with Tyrode’s buffer (12mM NaHCO_3,_ 10mM HEPES, 137 mM NaCl, 2.7mM KCl, 5.5mM glucose and 1 mM CaCl_2_/MgCl_2_, 1 volume blood: 4 volume Tyrode ratio). Diluted whole blood (50 μL) was stained at 37°C for 30 minutes with Antibodies and dyes listed in Supplementary Table 2. Thereafter, samples were diluted with Tyrode’s buffer (1 volume of blood: 10 volume of buffer ratio) before examination by flow cytometry. We characterized platelet activation as follows: CD41^+^-P-selectin^+^-Annexin V^+^. Mouse enoxaparin-anticoagulated whole blood was diluted with Tyrode’s buffer supplemented with 1 mM CaCl_2_/MgCl_2_ (1 volume blood: 2 volume Tyrode ratio) and treated similarly as human whole blood. For kinetics of activation in HOCl-SSc mice, whole blood was collected via sub-mandibular vein.

For each marker, MFI was obtained by background subtraction obtained from isotype control antibody or unstained samples when applicable. All samples were acquired with a BD LSRFortessa™ Cell Analyzer (BD biosciences and recorded data analyzed with FlowJo (v.10.10.0).

### Preparation of washed platelets and examination of activation profile by flow cytometry

Human or murine washed platelets were prepared as described (*93*) with slight modifications. Anti-coagulated blood was diluted with platelet washing buffer, PWB (4.3 mM Na_2_HPO, 24.3 mM NaH_2_PO, 4.3 mM K_2_HPO_4_, 113 mM NaCl, 5.5 mM glucose, 0.5% BSA, and 10 mM theophylline, pH 6.5, 1:1 ratio) containing enoxaparin (20 U/ml clexane) and apyrase (0.01 U/ml). Diluted platelet-rich plasma (PRP) was obtained by centrifugation at 37°C for 5 mins at 250g without break. Platelets in diluted PRP were then pelleted by centrifugation at 2000 x g for 5 mins at 37°C and washed twice in PWB and then finally resuspended (10^6^/mL) in Tyrode’s buffer (10 mM Hepes, 12 mM NaHCO_3_, 137 mM NaCl, 2.7 mM KCl, 5 mM glucose, pH 7.4) supplemented with 1 mM CaCl_2_/MgCl_2_ and 0.02 U/ml Apyrase. Platelets were let rest for 30 mins at 37°C, before stimulation and staining. We characterized P-selectin and Annexin V expression on singlet CD41+ population. All samples were acquired with a BD LSRFortessa™ Cell Analyzer (BD biosciences and recorded data analyzed with FlowJo (v.10.10.0).

### Whole lung homogenization

Whole lung samples were perfused and collected onto a 15mL falcon tube containing 500 µL of tissue protein extraction buffer (Pierce T-PER™, Tissue Protein Extraction Reagent, Thermo Fisher Scientific) containing protease (cOmplete™, EDTA-free Protease Inhibitor Cocktail) and phosphatase (PhosSTOP™) inhibitors. Samples were submitted to homogenization at maximum speed for 1 minutes on ice (Fisherbrand™ 150 Handheld Homogenizer). Thereafter, samples were transferred to a clean microcentrifuge tube and centrifuged at 16,000 × g for 10 mins at 4°C. Supernatant was collected to a new microcentrifuge tube and stored at −80°C until analysis.

### NETs formation and quantification *in vitro*

Freshly isolated neutrophils (1 x 10^6^) in RPMI containing 10% FBS were seeded onto a 24-well plate and stimulated with vehicle, A23187 (5 μM), or platelets (ratio 1 neutrophil: 50 platelets) for 3 or 4 h, human or mouse, respectively. Thereafter, each well was washed twice to removed unbound DNA before applying a solution of S7 nuclease (50 U/ml) in RPMI containing 10% FBS and incubation at 37°C for 30 minutes. The supernatant containing DNA and citrullinated H3 (citH3) was then transferred onto a microcentrifuge tube containing EDTA (5mM final) and centrifuged 5 minutes at 300g without break. The supernatant was then collected onto a clean microcentrifuge tube and stored at −80°C until analysis. Quantification of citH3 was performed on collected samples using the Citrullinated Histone H3 (Clone 11D3) ELISA Kit (Cayman Chemicals) according to manufacturer’s instructions. Absorbance at 450 nm was measured using a Synergy HTX multi-mode microplate reader.

### NET quantification *in vivo*

NETosis was determined as the quantity of citrullinated histone H3 (CitH3) detected in mouse sera (1:2 dilution) or lung tissue homogenates with the citrullinated histone H3 ELISA kit (Cayman Chemical).

### Neutrophil count in bronchoalveolar lavage and lung

Bronchoalveolar lavage (BAL) fluid was collected on euthanized mice via isoflurane overdose by gently washing the lungs with 1 ml sterile PBS for three times in the bronchial tree using a 22g endotracheal tube (Kent scientific). Lungs were then perfused at a rate of 200µL/second using a 3mL syringe with a 22g needle containing PBS into the right ventricle of the heart. Approximatively 2.5mL of PBS are generally used per lung. Whole lung samples were then harvested and submitted to enzymatic digestion in RPMI 1640 containing collagenase 2 (2mg/mL) and DNAse I (0.1 mg/mL) for 30 minutes at 37°C under constant mixing at 900 rpm in a thermomixer (Eppendorf® Thermomixer). After digestion, lung samples were submitted to a 70mM cell strainer on ice. Resulting cell suspensions from BAL and whole lung digestion were centrifuged at 500g for 5 mins at 4°C and resuspended with PBS/0.5% BSA before staining for neutrophils (anti-Ly6G antibody). Ly6G+ population was quantified by flow cytometry as percentage of all cells or absolute count using counting beads.

### Neutrophil depletion

Neutrophils were depleted using anti-mouse Ly6G (clone 1A8, BioXCell) injected in i.p at 100 μg for 7 consecutive days following by an injection every second day until the experiment end. For each experiment, rat IgG2a isotype control, anti-trinitrophenol (clone 2A3, BioXCell) was used as control. Neutrophil depletion efficiency was assessed by flow cytometry identified in the CD45^+^/Ly6C^+^ quadrant.

### Platelet depletion

For diphtheria toxin (DT)-induced platelet depletion, PF4^Cre+^/DTR^fl/fl^ and PF4^Cre-^/DTR^fl/fl^ (control) mice received i.p. injection of 250 ng of DT for 7 consecutive days following by an injection every three day until the experiment end. Platelet depletion in whole blood collected via the submandibular vein was assessed using Sysmex XP-300-Hematology-Analyzer.

### Adoptive transfer

Neutrophils were isolated from the bone marrow, T cell were isolated from the spleen, platelets were isolated from the blood. For platelet-activated neutrophils experiments, neutrophils were co-culture with platelets (ratio 1 neutrophil to 50 platelets) at 37°C for 3 h prior to treatment with trypsin/EDTA (0.01%) and centrifugation at 300g for 5 minutes to remove platelets prior neutrophil injection. Cells (5 x 10^6^) were injected i.d (skin fibrosis) or i.v. (lung fibrosis) into a recipient mouse. Tissues were collected after 14 days before tissue collection and processing as described above.

### Anti-GPVI mAb JAQ1 or irrelevant control IgG-treatment

Fibrosis was induced in WT mice by BLM injection as described above. On day 14 of the disease development, the anti-GPVI mAb JAQ1 or irrelevant control IgG were injected in the retro-orbital plexus (100 μg/mouse in sterile PBS) once a week. Mice were randomly allocated into groups. Between days 25 and day 28, mice were sacrificed for tissue harvest.

### Detection of Scl-70 in the serum

Anti–Scl-70 antibodies were detected in mouse sera (1:4 dilution) using a mouse anti-Scl-70 ELISA kit (Signosis) following manufacturer’s instruction. Absorbance at 450 nm was measured using a Synergy HTX multi-mode microplate reader.

### Total collagen content in lung

Total collagen content in lung was measured from whole lung homogenate using the total collagen assay kit (Quickzyme Biosciences). Results were expressed as the total collagen content in lung (mg/whole lung).

### Transcriptomic data

#### Acquisition of Transcriptomic Data

Publicly available transcriptomic datasets from SSc studies were obtained from the Gene Expression Omnibus (GEO), including datasets from whole blood (GSE130953 and GSE231691) and skin (GSE249550, GSE181549, GSE130955 and, GSE95065) and loaded into R (version 4.4.1) for analysis. Both microarray and RNA-seq experiments were employed. For data curation, only dSSc patients and healthy controls were included, employing baseline samples preferentially if multiple samples were available from a single dSSc patient. Microarray datasets were quantile-normalized and log2-transformed when needed using standard workflows in limma (version 3.60.6). Standard QC for RNA-seq raw counts were performed, low-expressed genes were removed using filterByExpr and normalization factors were estimated using the TMM (Trimmed Mean of M-values) method (edgeR version 4.2.2).

#### Differential expression analysis (DEA)

DEA was performed for each dataset according to its specifications. EdgeR-limma-voom pipeline was used for the RNA-seq analysis, modeling normalized log-transformed counts per million (CPMs) and accounting for batch and donor effect when needed. Linear models were used for microarray datasets with limma-voom (limma version 3.60.6). In all the DEA, SSc was used as the reference in the contrast, and genes that passed False Discovery Rate (FDR) < 0.5 were considered significant.

#### Gene Set Enrichment Analysis

Based on the Log2 Fold Change/Estimate statistics from the previous analysis, genes were ranked. For the analysis, fgsea package (version 1.30.0) was used, testing MSigDB C2 (package version 25.1.1), and using Reactome pathway terms. Terms passing 10% FDR were considered significant, and the Normalized Enrichment Score was used to identify the pathways enriched across datasets and their directions of change.

### Quantification and statistical analysis

Results were analyzed with GraphPad Prism statistical software (version 10.1.1). The statistical tests used are specified in the figure legends. Two-tailed unpaired t-tests were performed when comparing only two groups, Paired t-tests were used to compare internally controlled replicates, and ordinary one- and two-way ANOVA using Turkey’s multiple comparisons test was performed when comparing one variable across multiple groups. Sample sizes for each experiment are provided in the figures and the respective legends. P values are indicated in each figure.

## ACKNOWLEDGMENTS

R.D. was supported by an Arthritis National Research Foundation (ANRF) award, a National Scleroderma Foundation New Investigator Grant, and NIH/NIAMS P30 AR070253 pilot grants. L.S. was supported by NIH/NIAID T32 AI007512. M.G.A. was supported by NIH P30AR070253, the ARNF, the Vic Braden family, and the Lupus Research Alliance, the Charles H. Hood Foundation, and the Boston Children’s Hospital Office of Faculty Development/Basic & Clinical Translational Research Executive Committees Faculty Career Development Fellowship. P.A.J. was supported by PHOCEO AP-HM, Philippe Foundation, and Association France Vascularites. D.W. was supported by NIH/NHLBI R35HL166556. A.M.B. was supported by NIH/NHLBI 1R01HL155955. S.B.M. was supported by NIH/NHLBI K23HL15033 and R01HL171240 and reports research funding from Pliant Therapeutics and Boehringer Ingelheim. P.A.N. was supported by NIH/NIAMS 2R01AR065538, 2R01AR073201, the Rheumatology Research Foundation, the Lupus Research Alliance, and the Peabody Foundation.

---

**Supplemental figure 1.** Neutrophil migration in Human SSc. **(A)** Representative gating strategy for human peripheral neutrophils shown as the CD66b+ population gated on single cells. **(B)** Representative flow cytometry histogram plot for quantification of healthy control or dSSc neutrophil migration toward LTB4 or vehicle (no LTB4). **(C)** Right, neutrophil quantification in skin histology samples of controls (n=9) or SSc patients with skin fibrosis (n=17). Left, representative skin sections isolated from SSc patient showing the presence of MPO+ neutrophils (red arrowhead) in the dermis (original magnification 20x; MPO staining).

**Supplemental figure 2.** Transcriptomic dataset analysis. Reactome Neutrophil Degranulation Pathway Panel. Gene Set Enrichment Analysis (GSEA) plots for the Reactome *Neutrophil Degranulation* pathway across six SSc transcriptomic datasets. The running enrichment score curves (green) represent the cumulative enrichment of neutrophil-related genes throughout the ranked genome-wide list.

**Supplemental figure 3.** Neutrophil Gene Expression Heatmap. Differential expression of neutrophil-associated genes across six transcriptomic datasets in SSc. Colors represent log2 fold-change values estimated with edgeR (red = upregulated, blue = downregulated). Grey squares indicate genes not tested in each dataset. Statistical significance is indicated with stars (* adj. p < 0.05; ** < 0.01; *** < 0.001). Blood datasets displayed stronger neutrophil-related signaling compared with skin.

**Supplemental figure 4.** Mouse models of SSc. **(A)** Schematic representation of the HOCl-SSc model with timeline of injection for 42 days **(B).** Schematic representation of the BLM-SSc model with timeline of injection for 28 days.

**Supplemental figure 5.** Increased neutrophil infiltration in lung of HOCl and BLM mice **(A-C)** Representative flow cytometry histogram plot for Mac-1 (A) and L-selectin (B) expression and ROS generation (C) comparing PBS- (Blue) and HOCl (Red)-treated mice. (**D-E**) Neutrophils from PBS control and HOCl (**D**; n=11/group) or PBS control and BLM (**E**; n=6/group) were assayed for migration capacity toward the chemoattractant LTB4. Data are expressed as fold increased over no LTB4. Each dot is the average of duplicate results for each condition. The data represent the mean ± SD. **(F-K)** Quantification by flow cytometry of the neutrophil proportion and absolute count in the BAL **(F-G)** and in the lung **(H-K)** of PBS-, HOCl- or BLM-treated mice as listed (HOCl model: n = 10 PBS and n = 10 HOCl; BLM model: n= 9 PBS and n=9 BLM). The data represent the mean ± SD. BAL, bronchoalveolar lavage.

**Supplemental figure 6.** Neutrophil depletion and adoptive transfer. **(A)** Timeline representing the injection days of 1A8 or isotype control IgG (2A3) for neutrophil depletion in the HOCl model. **(B)** Representative flow cytometry histogram plots (left) showing neutrophil depletion in HOCl-treated mice after 49 days of 1A8 treatment compared to control IgG (n= 17/ group). Quantification of neutrophil proportion in the CD45+ leukocyte population is represented on the right.

**Supplemental figure 7.** Neutrophil and T cell adoptive transfer **(A)** Schematic representation of the adoptive transfer experiments. **(B)** Description of the flow-cytometric gating strategy to assess isolated neutrophil purity. **(C)** Description of the flow-cytometric gating strategy to assess isolated T cell purity**. (D)** Representative skin sections isolated from WT mice 2 weeks following adoptive transfer of T cells isolated from PBS- or HOCl-treated mice (original magnification 4x; Masson Trichrome staining). **(E)** Dermal thickness (right) was measured for each group (n=8 per group**). (F)** Representative lung sections isolated from WT mice 2 weeks following adoptive transfer of T cells isolated from PBS- or HOCl-treated mice (original magnification 20x; Masson Trichrome staining). **(G)** Microscopic scores of lung sections according to a modified Ashcroft’s method (right) were determined for each group (n=8 per group). The data represent the mean ± SD.

**Supplemental figure 8.** Neutrophils in PAD4-/- mice. **(A)** Schematic representing the different steps in the preparation of NETs samples for ELISA essay. **(B)** Levels of CitH3 in lung homogenates analyzed by ELISA comparing PBS (n=12) or SSc mice as listed in the figure (n= 11 HOCl; n= 10 BLM; n= 9 PAD4-/- HOCl; n=6 PAD4-/-BLM). **(C)** Total collagen content in the lung from WT or PAD4-/- mice treated with HOCl (n=10 per group). **(D-I)** Flow cytometry quantification of whole blood neutrophil Mac-1 and L-selectin expression, and ROS generation in WT or PAD4-/- mice treated with HOCl (**D, F & H**) or BLM **(E, G & I).** The data represent the mean ± SD.

**Supplemental figure 9.** Platelet Gene Expression Heatmap. Differential expression of platelet-associated genes across six transcriptomic datasets in SSc. Colors represent log2 fold-change values estimated with edgeR (red = upregulated, blue = downregulated). Statistical significance is indicated with stars (* adj. p < 0.05; ** < 0.01; *** < 0.001).

**Supplemental Figure 10.** Platelet depletion in SSc. **(A)** Description of the flow-cytometric gating strategy to assess whole blood platelets. **(B)** Representative flow cytometry histogram plot for showing the difference of P-selectin expression between groups. **(C)** Representative flow cytometry histogram plot for showing the difference of Phosphatidyl serine expression between groups. **(D)** Timeline representing the injection days of diphtheria toxin (DT) for platelet depletion in the HOCl model. **(E)** Platelet count was assessed using Sysmex in whole blood of HOCl PF4cre-iDTRfl/fl (+ Platelets) or HOCl PF4cre+-iDTRfl/fl (- Platelets) mice after 49 days of DT treatment (n=8/ group). The data represent the mean ± SD.

**Supplemental figure 11.** Feedback activation of platelets by neutrophils in SSc. Whole blood platelets from HOCl or BLM treated mice where neutrophils or NETs have been inhibited were assayed by flow cytometry. **(A - D)** Quantification at day 49 or 35 of P-selectin **(A & B)** and phosphatidylserine **(C & D)** on platelets in whole blood of HOCl- **(A & C)** or BLM- **(B & D)** treated mice, respectively, undergoing IgG or 1A8 treatment (HOCl model: n = 11 PBS and n = 11 HOCl; BLM model: n= 10 PBS and n=10 BLM). **(E - H)** Quantification at day 42 or 28 of the levels of P-selectin **(E & F)** and phosphatidylserine **(G & H)** on platelets in whole blood of WT or PAD4-/- mice treated with HOCl **(E & G)** or BLM (**F & H**), respectively (HOCl model: n = 15 PBS and n = 15 HOCl; BLM model: n= 14 PBS and n=14 BLM). The data represent the mean ± SD.

**Supplemental figure 12.** Platelets are more susceptible to collagen stimulation in SSc. **(A & B)** Representative flow cytometry histogram plots (Top) showing the difference in P-selectin **(A)** phosphatidylserine **(B)** expression to determine platelet reactivity toward various agonist (as listed) between healthy control (HC) and dSSc patients (n=3 per group). Quantification for each surface markers are represented below the histogram for each agonist. **(C & D)** Quantification of platelet reactivity towards various agonist in mouse platelets isolated from PBS- or HOCl-treated mice (n=5/group). **(E & F)** Flow cytometry quantification of GPVI receptor on surface of platelets from GPVI^+/+^ or GPVI^-/-^ mice treated with HOCl **(E)** or BLM (**F**; HOCl model: n = 10 PBS and n = 12 HOCl; BLM model: n= 15 PBS and n=14 BLM). The data represent the mean ± SD.

**Supplemental figure 13. (A-D)** Whole blood platelets and neutrophils were assayed by flow cytometry in GPVI^-/-^ or littermate control GPVI^+/+^ mice treated with HOCl at d42 (n = 13 PBS and n = 15 HOCl). Platelets were assayed for P-selectin and phosphatidyl serine expression in the HOCl model **(A & B)**. Neutrophils were assayed for Mac-1 and L-selectin expression in the HOCl model **(C & D). (E)** Levels of CitH3 in serum of GPVI^-/-^ or WT mice treated with HOCl analyzed by ELISA (n=10 per group). (**F**) Representative skin sections isolated from GPVI^+/+^ or GPVI^-/-^ mice treated with HOCl (original magnification 4x; Masson Trichrome staining). **(G)** Dermal thickness was measured for each group (n = 13 PBS and n = 15 HOCl). **(H)** Representative lung sections isolated from GPVI^+/+^ or GPVI^-/-^ mice treated with HOCl (original magnification 20x; Masson Trichrome staining). **(I)** Microscopic scores of lung sections according to a modified Ashcroft’s method (right) were determined for each group (n = 13 PBS and n = 15 HOCl). (**J**) Flow cytometry quantification of GPVI receptor on surface of platelets PBS- or BLM-mice treated with JAQ1 or IgG. (**K & L**) Whole blood platelets were assayed by flow cytometry in PBS or BLM-mice treated with JAQ1 or IgG at d28 (PBS: n=10, IgG: n= 10, JAQ1: n=9). Platelets were assayed for P-selectin (**K**) and phosphatidyl serine expression (**L**). (**M**) Western blot quantification of α-SMA in lung lysates of BLM-mice treated with JAQ1 or IgG.

