## Supplemental files for "A GPVI-platelet-neutrophil-NET axis drives systemic sclerosis"

### Supplemental Figure 1: Neutrophil migration in Human SSc

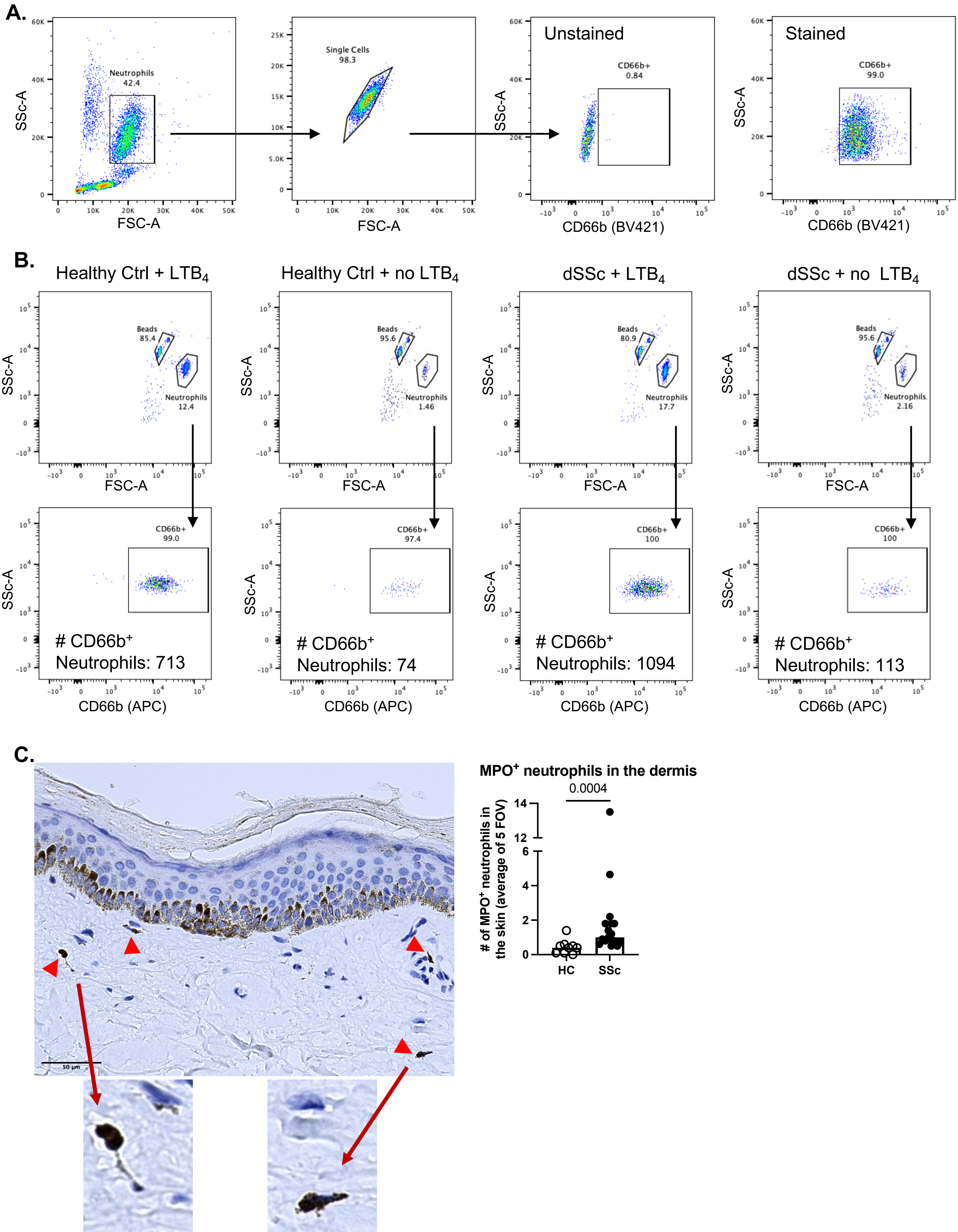

### Supplemental Figure 2: Neutrophil degranulation pathway in SSc

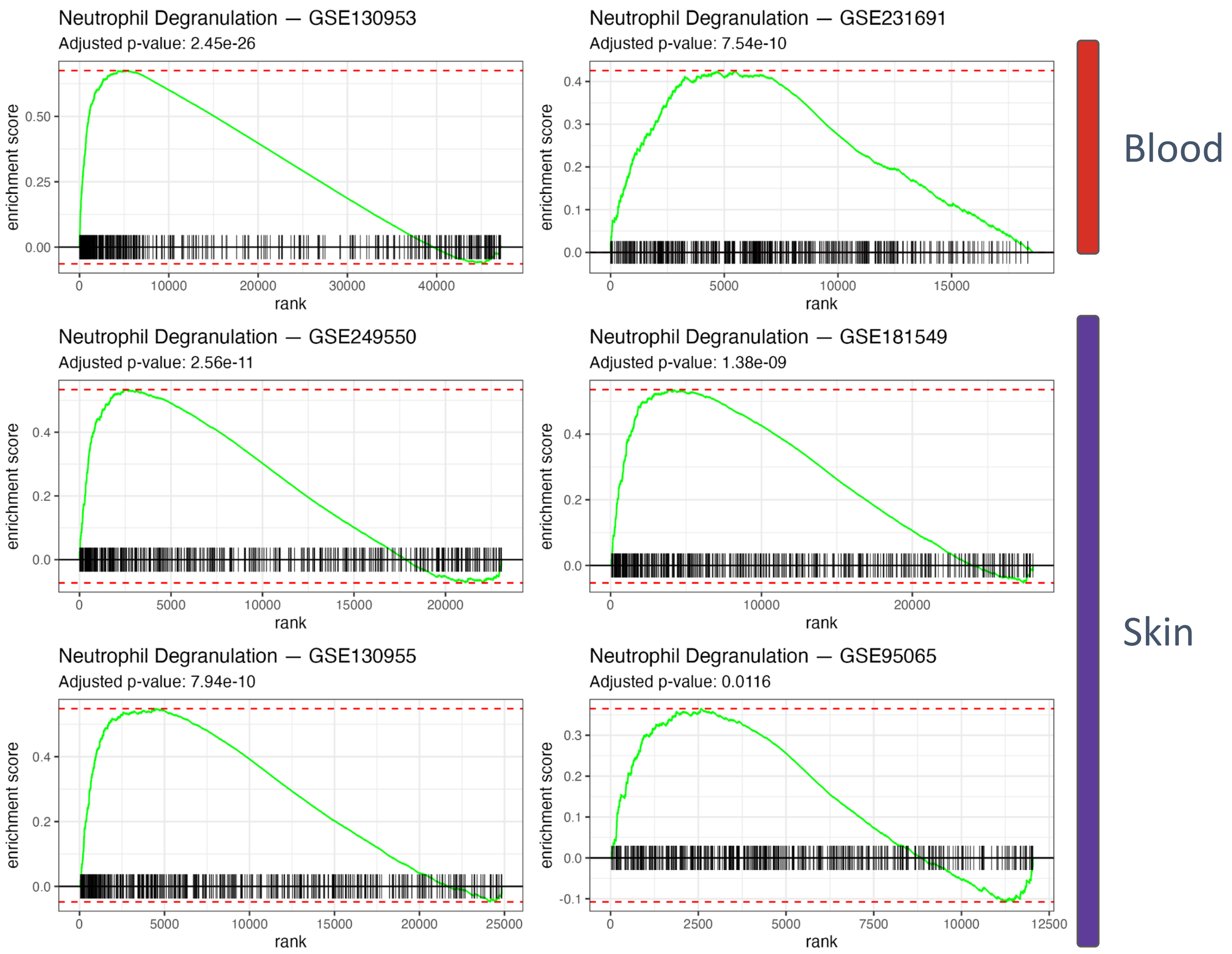

**Supplemental Figure 3: Neutrophil Gene Expression Heatmap.**

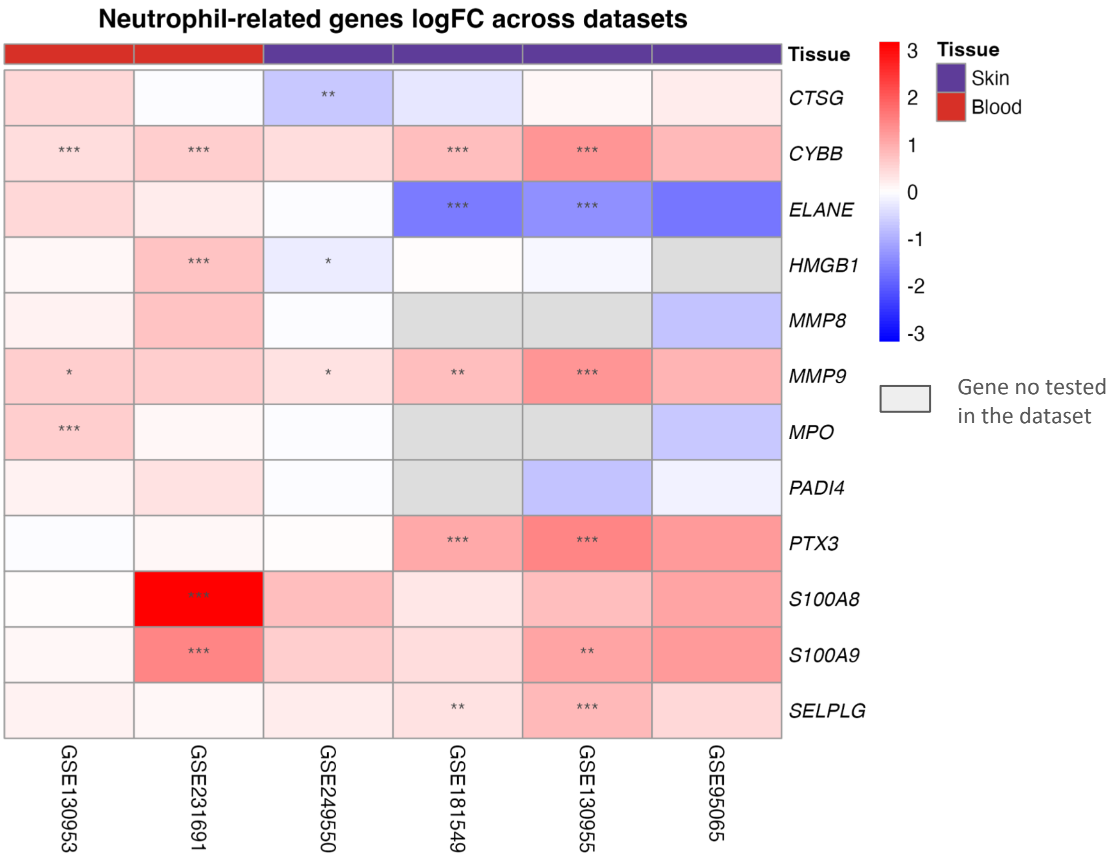


*: padj < 0.05; **: padj < 0.01; *** : padj < 0.001

#### Supplemental Figure 4: Mouse models of SSc

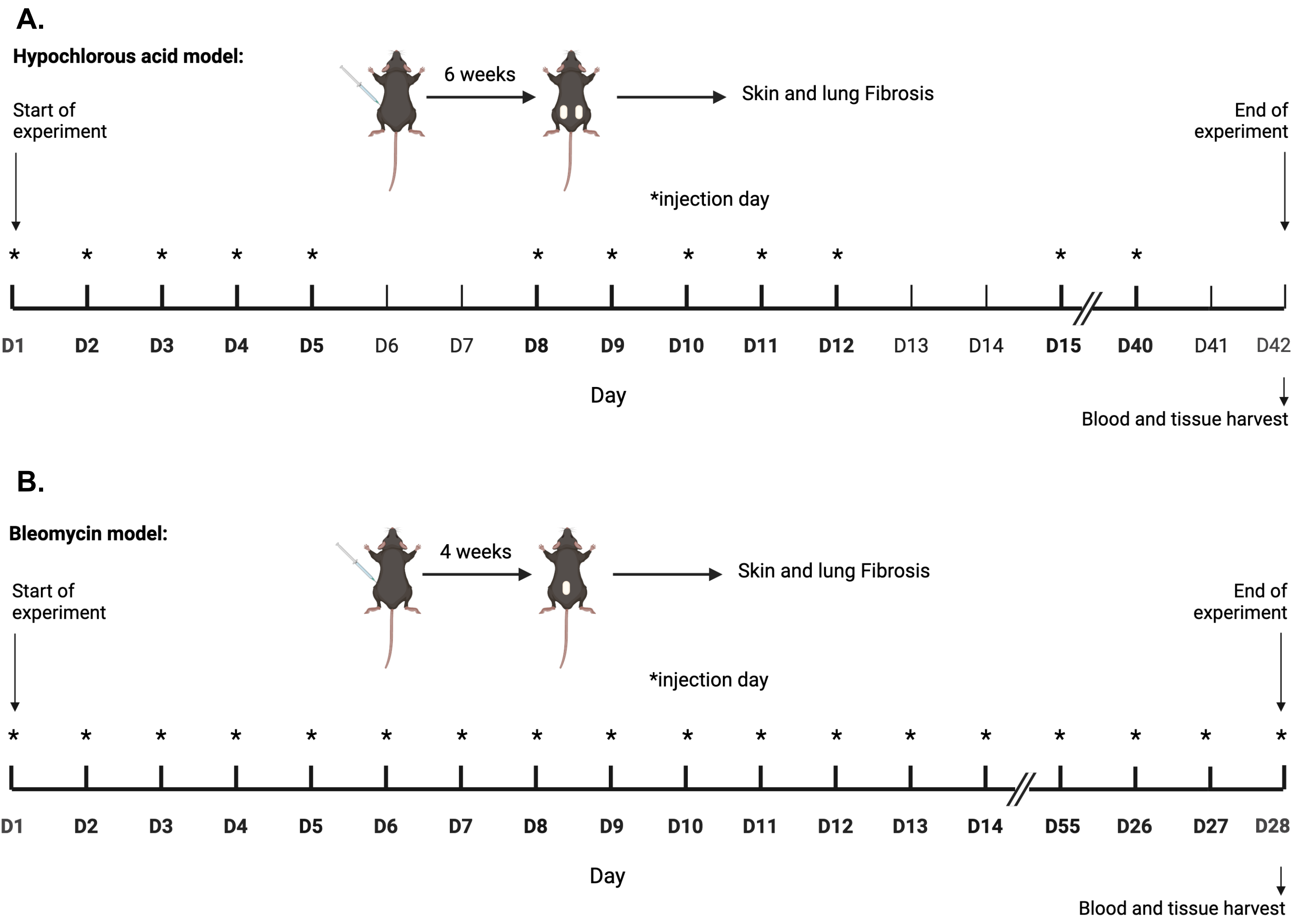

**Supplemental Figure 5: Increased neutrophil infiltration in lung of HOCl and BLM mice**

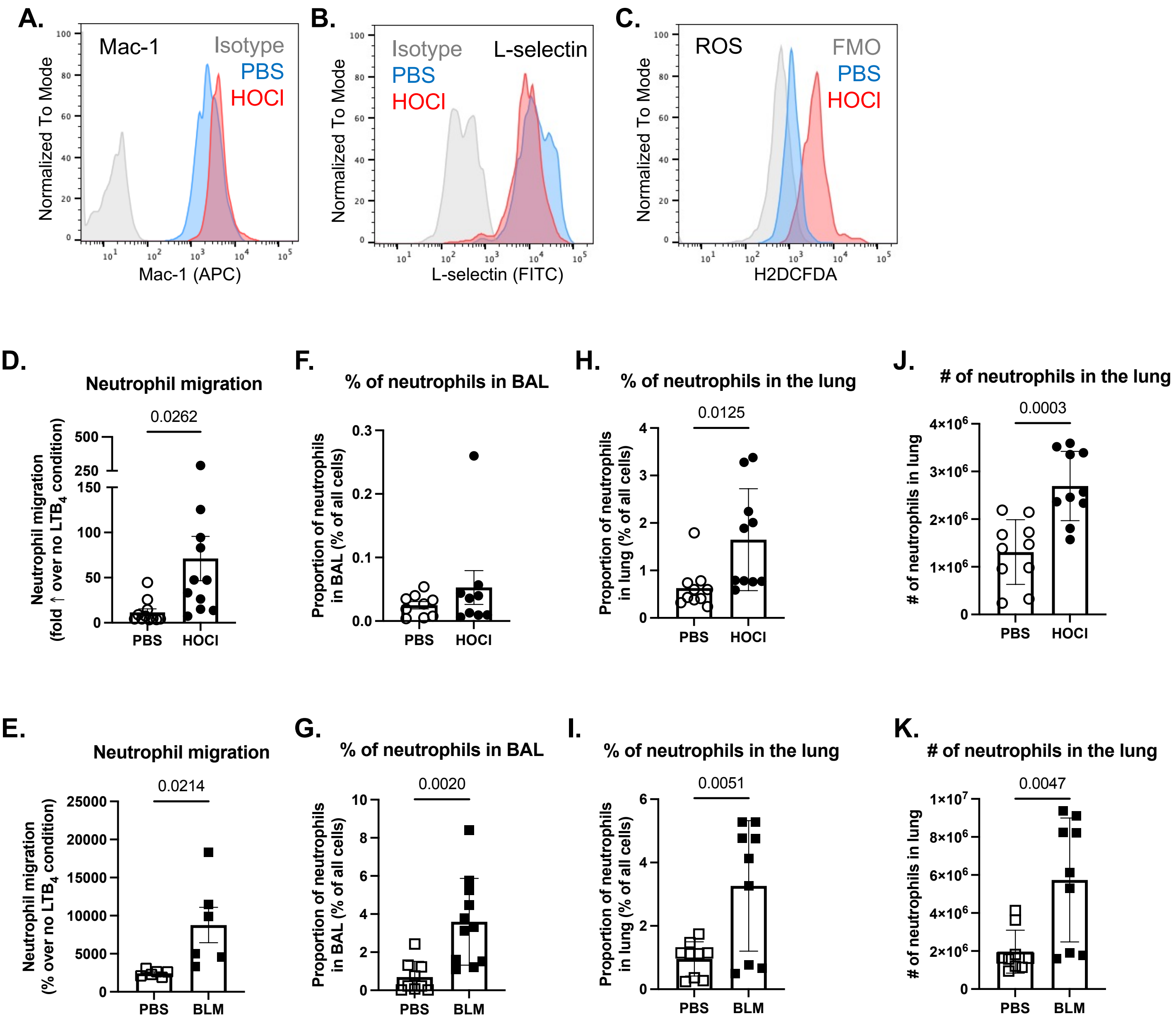

### Supplemental Figure 6: Neutrophil depletion

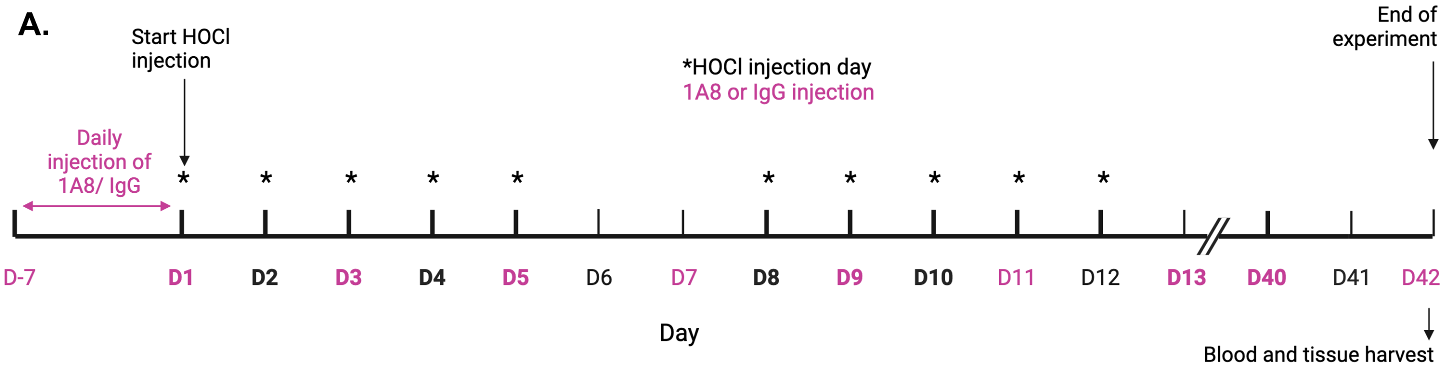


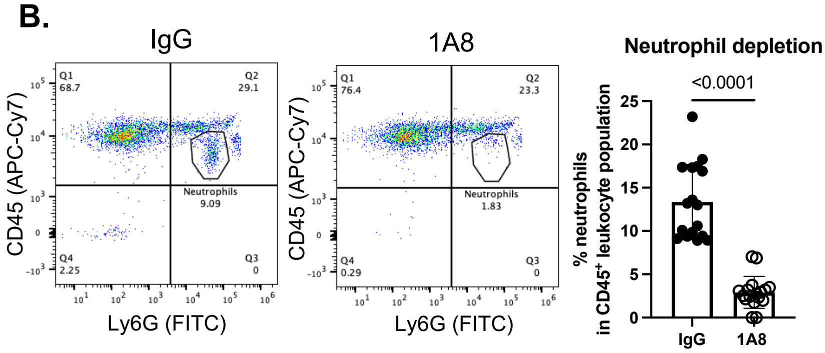

Supplemental Figure 7: Neutrophil and T cell adoptive transfer

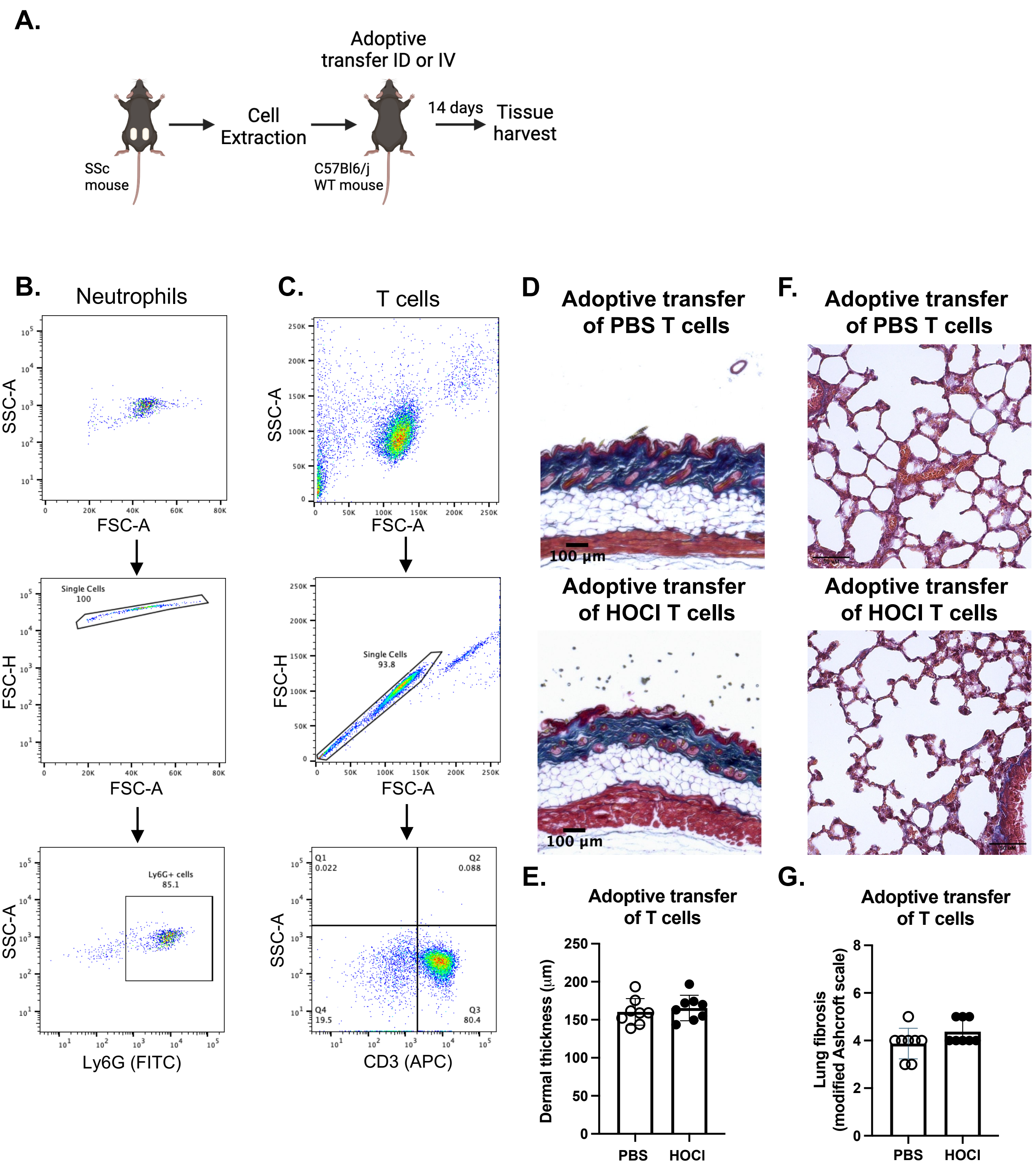

### Supplemental Figure 8: Neutrophils in PAD4$^{-/-}$ mice

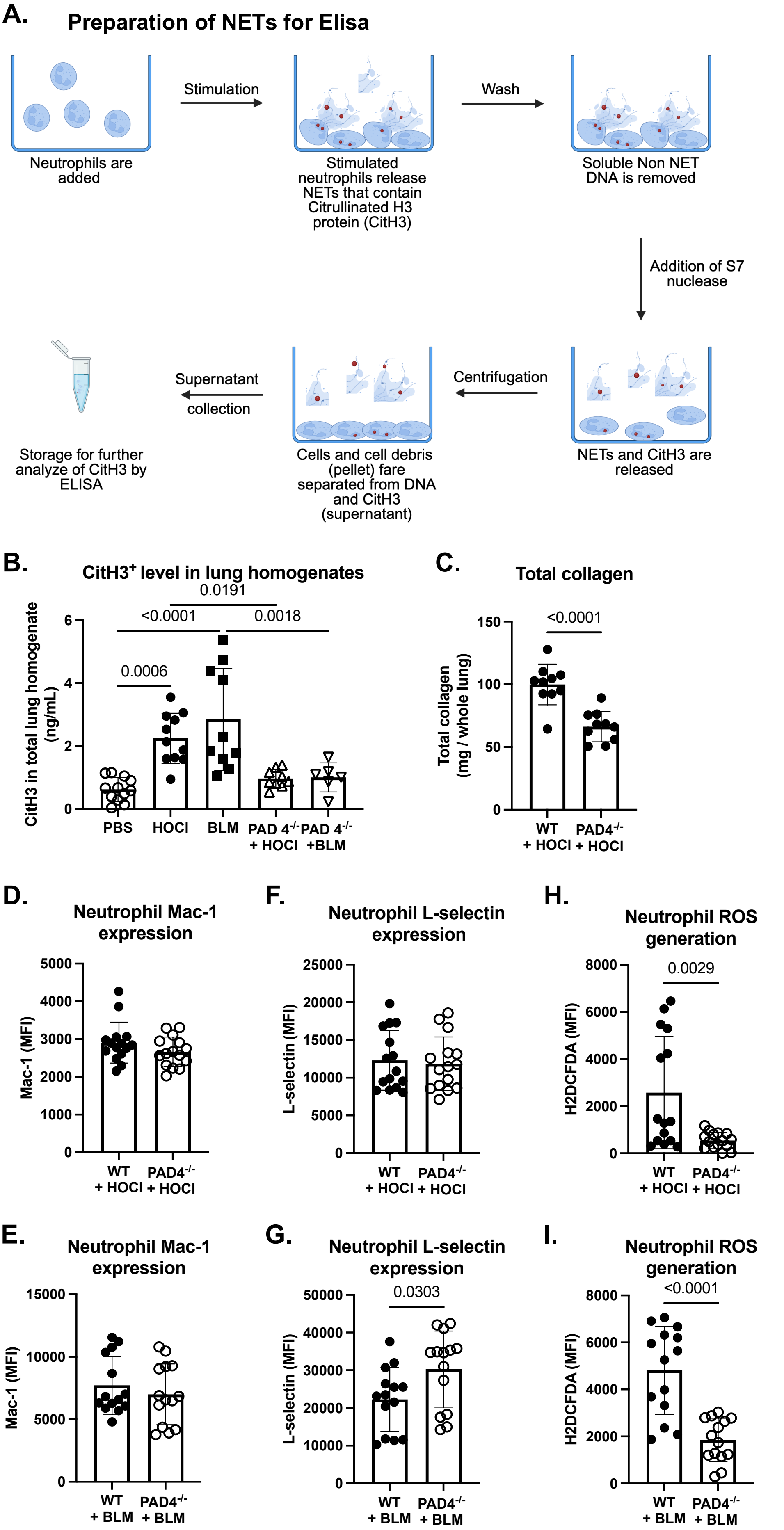

**Supplemental Figure 9: Platelet Gene Expression Heatmap**

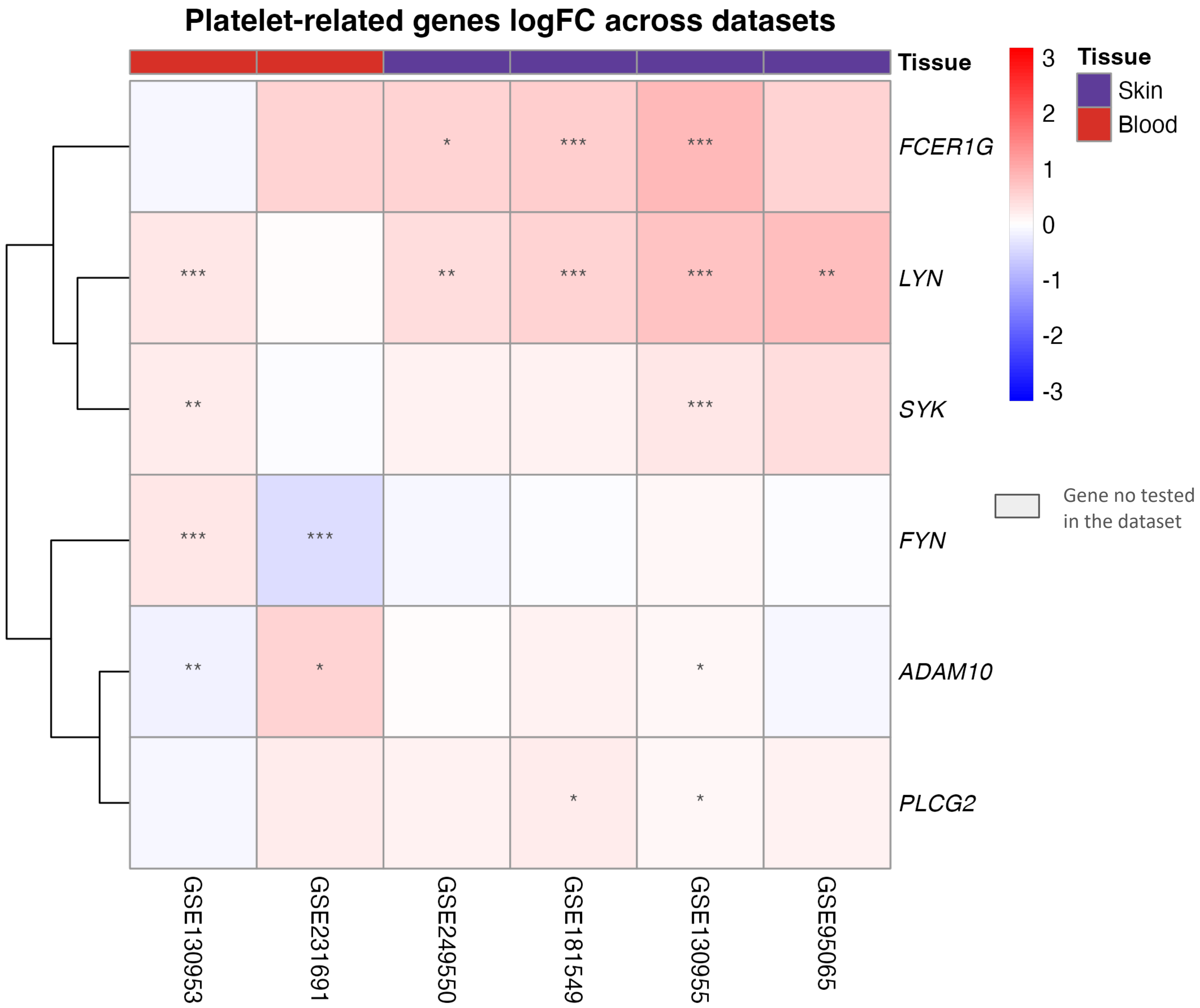


*: padj < 0.05; **: padj < 0.01; *** : padj < 0.001

#### Supplemental Figure 10: Platelet depletion in SSc

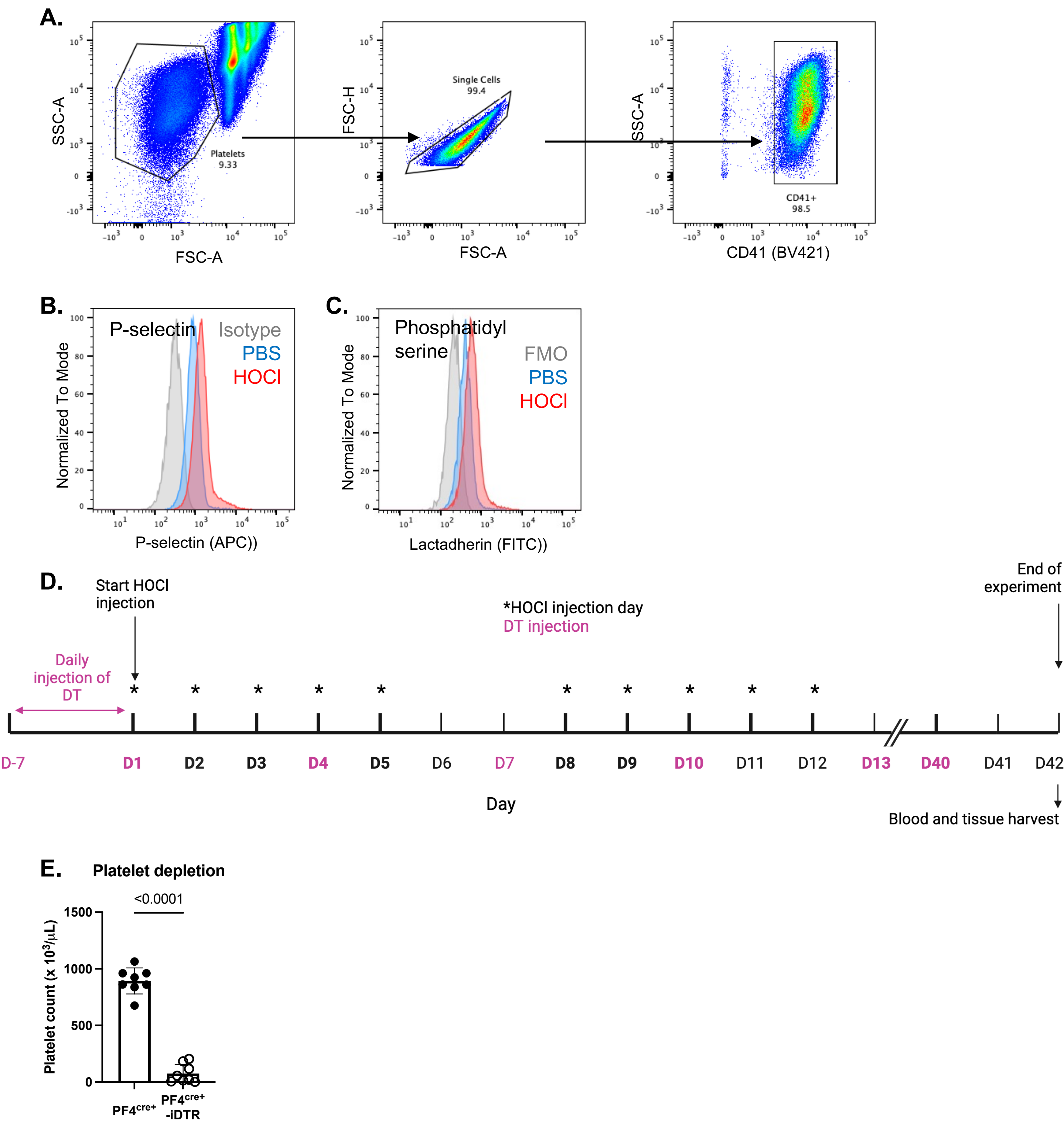

### Supplemental Figure 11: Feedback activation of platelets by neutrophils in SSc

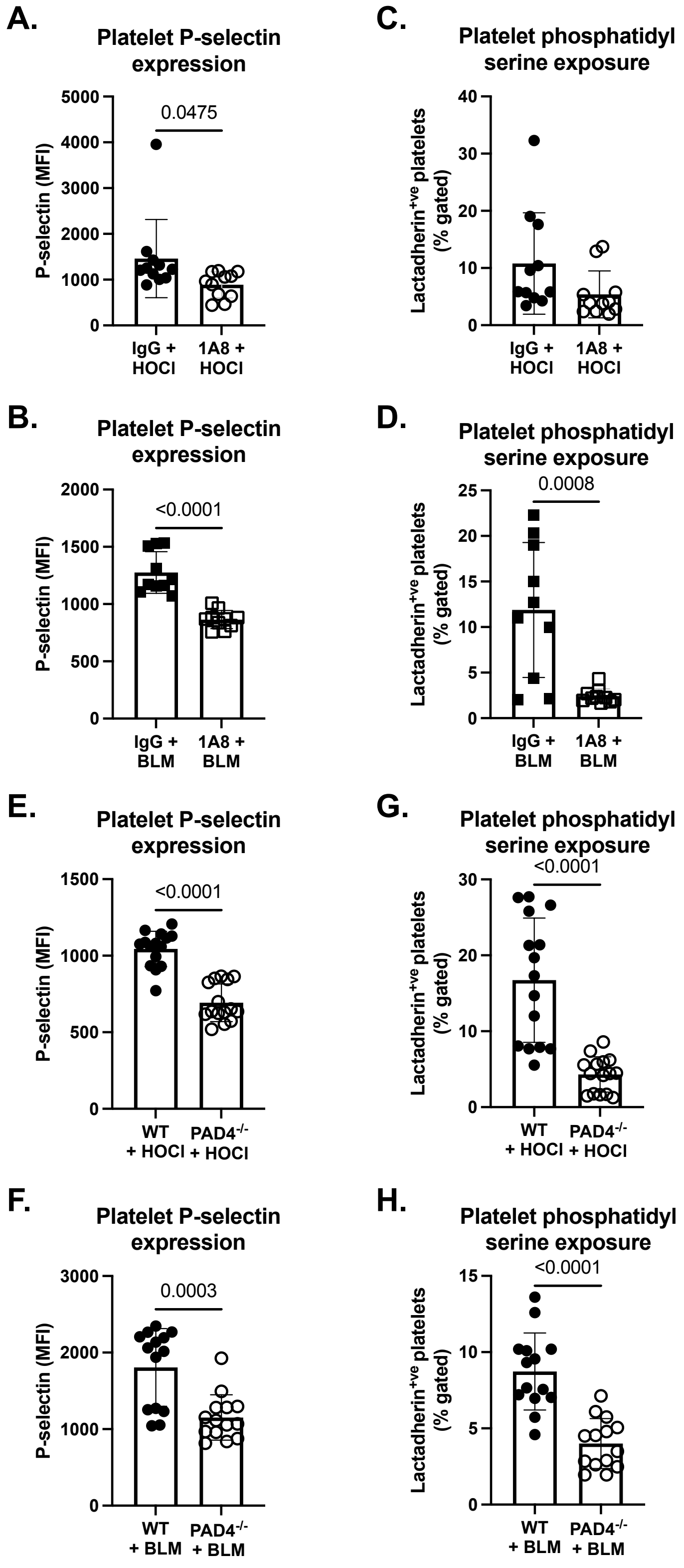

### Supplemental Figure 12: Platelets are more susceptible to collagen stimulation in SSc

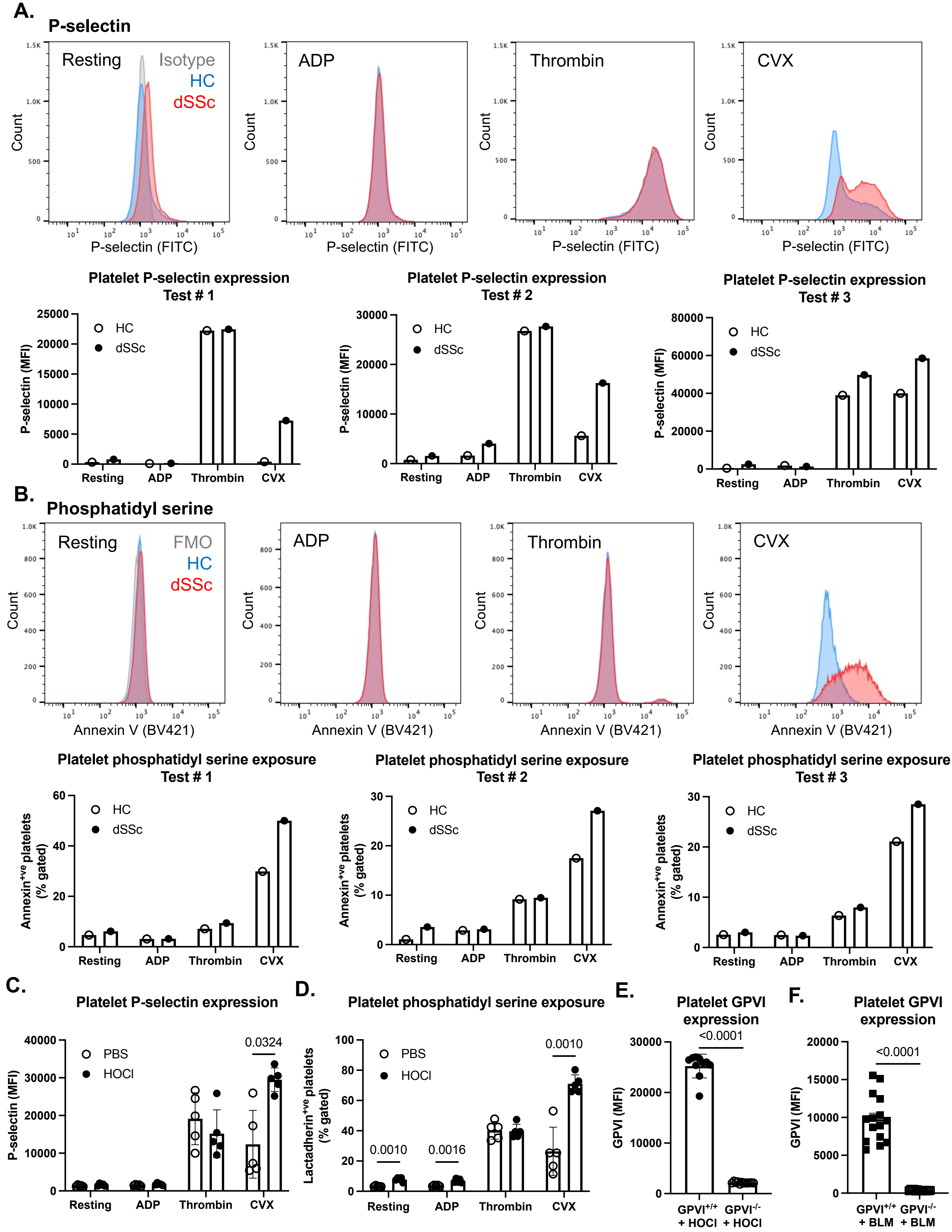

#### Supplemental Figure 13:

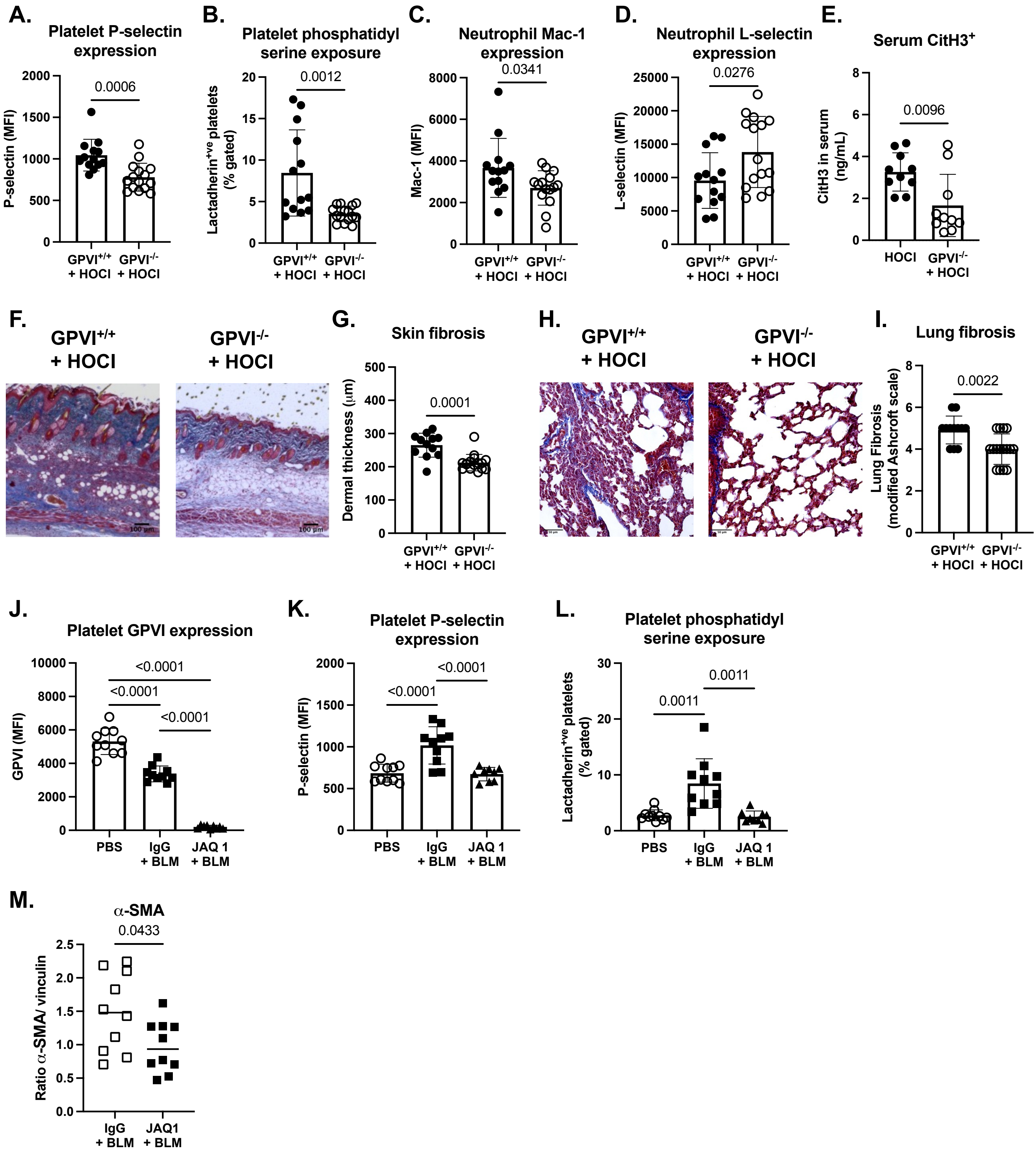

#### Supplemental Table 1: Demographic and phenotypic information of dSSc patients

| Patient | Age (years) | Sex | Disease | Complications/Co-morbidities | Medications | Other Auto-antibodies |
|---|---|---|---|---|---|---|
| 1 | 35 | Female | Diffuse SSc | Raynaud's | MMF, sildenafil | Scl-70 + , RNP +, ANA 1:1280 |
| 2 | 43 | Female | Diffuse SSc | ILD, depression | MMF, aripiprazole, nifedipine, venlafaxine | Scl-70 n/a |
| 3 | 62 | Female | Diffuse SSc | ILD, Sjogren's | MMF, omeprazole | Scl-70 -, RNP (equivocal) +, RNA Pol 3 +, SS-A +, SS-B + |
| 4 | 69 | Male | Diffuse SSc | ILD / restrictive lung disease, PH, Raynaud's | MMF, tadalafil | Scl-70 +, ANA 1:640, IgG + |
| 5 | 66 | Female | Diffuse SSc | ILD, Raynaud's | Botulinum toxin A injections | Scl-70 -, RNA Pol 3 +, ANA 1:640 |
| 6 | 60 | Male | Diffuse SSc | ILD, Raynaud's | MMF, Botulinum toxin A injections | Scl-70 +, ANA 1:2560, Pm/SCL + |
| 7 | 55 | Male | Diffuse SSc | Renal crisis, HTN, T2DM, Prior alcoholism, Raynaud's | MMF, amlodipine | Scl-70 -, RNA Pol 3 +, ANA 1:640, cardiolipin IgG + |
| 8 | 69 | Male | Diffuse SSc | ILD, esophageal dysmotility, GERD | MMF | Scl-70 -, ANA 1:1280, Th/To + |
| 9 | 51 | Male | Diffuse SSc | ILD, Arthritis | MMF, leflunomide | Scl-70 -, ANA 1:2560 |
| 10 | 56 | Female | Diffuse SSc | ILD in the context of calcinosis, telangiectasia | None | Scl-70 +, ANA - |
| 11 | 75 | Male | Diffuse SSc | ILD, Cardiomyopathy, GERD, Raynaud's | None | Scl-70 +, ANA 1:1280, Ro60 + |
| 12 | 36 | Female | Diffuse SSc | ILD | MMF | Scl-70 -, PM-Scl + |
| 13 | 77 | Female | Diffuse SSc | PH, Osteoporosis, Raynaud's | Treprostinil, HCQ, prednisone, rituximab, metoprolol | Scl-70 -, ANA 1:2560, Anti-Ro60 +, Th/To + |

SSc: systemic sclerosis; ILD: interstitial lung disease; PH: pulmonary hypertension; HTN; T2dM: type 2 diabetes mellitus; GERD: gastroesophageal reflux disease; MMF: mycophenolate mofetil; HCQ: hydroxychloroquine; RNP: ribonucleoprotein; ANA: antinuclear antibody; RNA Pol 3: RNA polymerase III

#### Supplemental Table 2: Demographic and phenotypic information of healthy controls

| Healthy Control | Age | Gender | Medications |
|---|---|---|---|
| 1 | 33 | Female | Hormonal birth control (pill) |
| 2 | 26 | Female | n/a |
| 3 | 39 | Female | n/a |
| 4 | 35 | Male | n/a |
| 5 | 37 | Female | Hormonal birth control (IUD), topamax (migraine) |
| 6 | 40 | Male | n/a |
| 7 | 39 | Male | n/a |
| 8 | 40 | Male | n/a |
| 9 | 33 | Male | n/a |
| 10 | 24 | Female | n/a |
| 11 | 39 | Male | n/a |
| 12 | 28 | Female | n/a |
| 13 | 38 | Female | n/a |

### Supplemental Table 3: Demographic and phenotypic information of skin biopsy donors

| Patient | Age | Gender | mRSS | Complications/Co-morbidities | Medications | Other Auto-antibodies |
|---|---|---|---|---|---|---|
| 1 | 34 | Female | 14 | Dyspnea, cardiac fibrosis, skin sclerosis, hypomotility, intestinal psuedoobstruction, Raynauds, Inflammatory arthritis , hypertension, proteinuria | IVIG, lasix, lisinopril | ANA + 1;1280, nucleolar; CCP + |
| 2 | 34 | Female | | | | |
| 3 | 26 | Female | 24 | Gastroparesis, dyspnea, Raynaud's | Cellcept, prednisone, nifedipine, IVIG | ANA + 1:1280; nucleolar pattern; centromere+ (2.6); PL-7 (27); OJ (15); K6 |
| 4 | 27 | Female | | | | |
| 5 | 59 | Male | 23 | Raynaud's, telangiectasias, hypertension, coronary artery disease, hyperlipidemia, alcohol abuse | CellCept, Nifedipine | RNA pol 3+; ANA+ 1;320 ,;ANA+ 1:1280 ; RNP + |
| 6 | 38 | Female | 21 | GERD, Raynaud's, dyspnea, Ehigh blood pressure, pulmonary nodules | Mycophenolate | ANA+; RNA pol 3+ |
| 7 | 38 | Female | | | | |
| 8 | 58 | Female | 18 | Raynaud's, fingertip lesions, Abnormal nailfold capillaries, dermatiis | Prednisone, gabapentine, cyproheptadine, halobetasol , Pimecrolimus, Calcipotriene, Tacrolimus | |
| 9 | 52 | Male | 10 | Raynaud's , ILD, SLE | Myfortic, MMF | ANA+, Scl70+ |
| 10 | 56 | Female | 2 | Raynaud's, GERD, sclerodactyly, Hashimoto's thyroiditits | Gabapentin, Plaquenil, methotrexate | Scl-70+; SRP Ab- 12 H; MI-2 Alpha Ab- 48 H; ANA + |
| 11 | 49 | Female | | | | |
| 12 | 63 | Female | 12 | GERD | Azathioprine (Imuran), Amlodipine, Lisinopril | ANA+ |
| 13 | 63 | Female | | | | |
| 14 | 46 | Female | 6 | ILD, arthritis, heart failure, thyroid disease | Prednisone, Amlodipine, Losartan | ANA+ |
| 15 | 45 | Female | | | | |
| 16 | 60 | Male | 25 | GERD, pulmonary hypertension, heart failure, arthritis, obesity, thyroid disease | N/A | RNA pol 3 + |
| 17 | 57 | Male | | ILD, arthritis, anemia, pancytopenia, anxiety, AIHA | Prednisone, tadalafil (Cialis), | Scl-70 + |
| 18 | 44 | Male | | | | |
| 19 | 26 | Female | 7 | Telangiectasia | Smethicone | N/A |
| 20 | 67 | Male | 42 | Raynaud's, arterial fibrilation | Cellcept, prednisone, | N/A |
| 21 | 71 | Male | | | | |
| 22 | 34 | Female | 27 | Toxic maculopathy of retina, heart failure, pericarditis | IV Cyclophosphamide, Prednisone, Plaquenil, Gabapentin, nifedipine, amitryiptyline, | ANA + |
| 23 | 41 | Female | 24 | GERD | Rilonacept trial | N/A |
| 24 | 43 | Female | 21 | Dyspnea, sclerodactyly, kin sclerosis, irregular heart rhythm | IV Cytoxan, bactrim Ds, omeprazole, prochlorperazine | n/a |
| 25 | 41 | Female | | | | |
| 26 | 36 | Female | 22 | N/A | Cellcept, prednisone | N/A |
| 27 | 31 | Female | | | | ANA +, Anti-RO 60 +, Th/To + |

#### Supplemental Table 4: Transcriptomic datasets

| GEO | Organ | Assay | dSSc | Controls | Observation | PMID |
|---|---|---|---|---|---|---|
| GSE130953 | Blood | Microarray | 72 | 44 | Only dcSSc samples, baseline visit | 31391177 |
| GSE231691 | Blood | RNA-seq | 49 | 18 | -- | 37867940 |
| GSE130955 | Skin | RNA-seq | 48 | 33 | Only baseline biopsy | 31767698 |
| GSE181549 | Skin | Microarray | 70 | 44 | Only dSSc samples at baseline | 21360508 |
| GSE249550 | Skin | Microarray | 48 | 33 | Only baseline biopsy | 38272035 |
| GSE95065 | Skin | Microarray | 18 | 15 | Two technical batches | -- |
